# Dynamic TET1 repression coordinates activation of macroH2A.2-poised bivalent genes during neuroectoderm differentiation

**DOI:** 10.64898/2026.10.17.759687

**Authors:** Brendan Terry, Carine Stapel, Fatima Santos, Joshua Kent, Kamal Kishore, Clive D’Santos, Ian Chambers, Aled Parry, Wolf Reik

## Abstract

Cell state transitions involve changes in gene expression programs in conjunction with epigenetic regulation occurring in chromatin. Bivalent chromatin, marked by overlapping active H3K4me3 and repressive H3K27me3, poises key developmental genes during pluripotency for cell type-specific activation during differentiation. However, the molecular mechanisms that activate bivalent genes during lineage commitment remain poorly understood. Here, we elucidate how the cytosine dioxygenase TET1 promotes neuroectoderm differentiation through dynamic chromatin binding and a non-catalytic repressor function. Through a hypothesis-free screen for TET1 chromatin complex partners, we find that TET1 physically interacts with macroH2A.2 and H2A.X on chromatin. Integrated multi-omic analyses reveal that the non-catalytic repressor function of TET1 and a novel activating function of the H2A variant macroH2A.2 together promote a poised transcriptional state within bivalent chromatin during pluripotency. Upon pluripotency exit, TET1 disengages from macroH2A.2-marked early neuroectoderm genes to permit their activation, while remaining bound as a repressor at mesoendoderm and late neural genes to prevent alternative cell state transitions. Furthermore, TET1 retention at accessible primed enhancers supports neural program activation. Together, our findings reveal a noncanonical functional interplay within bivalent chromatin between TET1 and macroH2A.2 that maintains a poised state during pluripotency and predicts selective gene activation during neuroectoderm specification.

## INTRODUCTION

A complex interplay between cell-extrinsic signalling and cell-intrinsic regulation of gene expression by transcription factors and the epigenetic landscape directs cellular differentiation during mammalian development (1–6). During gastrulation, the pluripotent epiblast gives rise to the primary germ layers of endoderm, ectoderm, and mesoderm, as well as their derivative early organogenesis cell types (7, 8). The pluripotent epiblast undergoes an Epithelial-Mesenchymal Transition (EMT) and migration through the Primitive Streak (PS) to form mesoderm and endoderm (9–12). In contrast, neuroectoderm derives directly from epiblast cells (13–16). Mouse and human pluripotent stem cells exhibit a default lineage bias towards neuroectoderm in the absence of external inductive signals, indicating an intrinsically encoded developmental hierarchy (1, 17–20). *In vivo* neuroectoderm specification depends on the cell-intrinsic epigenetic regulation by the TET family of cytosine dioxygenases, whose catalytic activity underlies active DNA demethylation (21, 22). Within the TET enzyme family, TET1 is uniquely expressed from pre-implantation through mid-gastrulation and drives the naive-to-primed pluripotency transition and neuroectoderm formation (23–27). Loss of full-length TET1 impairs neural progenitor cell (NPC) specification, increases mesoendodermal competence through dysregulation of Wnt and Nodal signaling, and causes gastrulation failure (24, 28). Importantly, catalytic mutants of TET1 target chromatin normally, complete gastrulation, and retain neuroectoderm differentiation competence, demonstrating that the non-catalytic function of TET1 is sufficient for neural fate specification during early development (24, 29–31).

Key developmental gene promoters in pluripotent cells are regulated by a dynamic bivalent chromatin state, in which enzymatic maintenance of activating H3K4me3 and repressive H3K27me3 balances gene silencing with the competence to respond to developmental cues (6, 32–42). During pluripotency, TET1 represses bivalent promoters non-catalytically by recruiting co-repressor complexes, including Polycomb Repressive Complex 2 (PRC2) and SIN3A, which preserve H3K27me3 levels (29, 43–45). Paradoxically, TET1 expression is also required for bivalency resolution—the loss of H3K27me3 and gain of H3K4me3—and corresponding gene activation during neuroectoderm differentiation (6, 46–49). How TET1 shifts from maintaining global bivalent silencing during pluripotency to enabling selective bivalent gene activation upon neuroectodermal fate commitment has remained unexplored.

Here, we demonstrate that TET1 non-catalytically coordinates with specific histone variants to control neuroectoderm commitment upon pluripotency exit. Through quantitative proteomic and chromatin interactome profiling during NPC differentiation from pluripotency, we identify macroH2A.2 and H2A.X as chromatin complex partners of TET1. Integrated ChIP-seq and transcriptomic analyses show that a TET1 and macroH2A.2 functional antagonism supports a poised transcriptional state at bivalent promoters during pluripotency. Upon pluripotency exit, TET1 selective disengagement from macroH2A.2-marked early neuroectodermal promoters permits gene activation, whereas persistent TET1 binding at mesoendodermal and late neuroectodermal promoters maintains silencing. Thus, TET1 regulates bivalent genes during neuroectoderm commitment through a mechanism of dynamic chromatin repression.

## MATERIALS AND METHODS

### ES cell culture

E14 mouse embryonic stem cells (ESCs) were purchased from the Cambridge Stem Cell Institute (wild-type) or were a generous gift of Prof. Ian Chambers (“TAG”) and cultured under serum/LIF conditions as previously described (50). TAG cells contain an endogenously modified *Tet1* leading to expression of TET1 with a C-terminal 3× FLAG tag, which have been characterised previously (26). Briefly, feeder-free ESCs were grown at 37 °C on gelatin-coated plates in normal oxygen under standard serum/LIF culture conditions (DMEM (Gibco, 11995-040), 2 mM GlutaMAX (Gibco, 35050-061), 15% fetal bovine serum (ThermoFisher Scientific), 1 U/mL penicillin – 1 mg/mL streptomycin (Gibco, 15140-122), 0.1 mM nonessential amino acids (Gibco, 11140-050), 50 mM β-mercaptoethanol (Gibco, 31350-010), 10^3^ U/mL LIF Stem Cell Institute, Cambridge)). Cell cultures were tested for mycoplasma contamination regularly (MycoAlert Mycoplasma Detection Kit, Lonza LT07–218) and always tested negative.

### Neural progenitor cell differentiation

After at least three passages in serum/LIF conditions, E14 ESCs were differentiated into neural progenitor cells (NPCs) using an optimised version of a published monolayer differentiation protocol (51, 52). Briefly, cells were seeded at 2,500 cells/cm² and the following day (Day 0) were transferred to modified N2B27 medium supplemented with insulin and BSA Fraction V. At Day 1, 1 μM all-*trans*-retinoic acid (RA) was added. These conditions were optimised in pilot experiments comparing seeding densities and RA treatment based on NPC differentiation efficiency assessed by immunofluorescence and qRT-PCR. For time-course experiments, biological replicates separated by ≥10 passages were collected at Days 0, 2, 4, and 6, washed with PBS, and lysed in 350–600 μL RLT buffer (Qiagen) for RNA extraction.

### HEK293T cell culture

HEK293T maintenance culture was performed using D10 media: (DMEM (Gibco,11995-040), 15% fetal bovine serum (Thermo Fisher Scientific) and 1 U/mL penicillin - 1 mg/mL streptomycin (Gibco, 15140-122)). Briefly, HEK293T cells were maintained in T175 flasks at a seeding density of 6 million cells in 30–40 mL D10 with passaging every 2–3 days (∼70% confluence). Lentiviral production in HEK293T cells was performed using ESC cell culture media without LIF.

### sgRNA plasmid generation

The top four sgRNA protospacers predicted by CRISPick were synthesized as oligos (Merck) with the following sequence design: 5’-TATCTTGTGGAAAGGACGAAACACCG-sgRNA protospacer-GTTTAAGAGCTAGGCCAACATGAGGATCACCCATG-3’ (53, 54). The individual oligos were cloned into the backbone CROP-sgRNA-MS2 (Addgene #153457) at BsmBI restriction sites using the NEBridge® Golden Gate Assembly Kit (BsmBI-v2, NEB) following manufacturer instructions. Constructs were validated by Sanger sequencing.

### Lentiviral production

Lentivirus was produced as described previously (50). Briefly, 3.5 × 10^6^ HEK293T cells were seeded in 10-cm dishes 16–20 h before transfection. For each sgRNA construct, 10 μg sgRNA plasmid was combined with 3.5 μg pMD2.G (Addgene #12259) and 6.5 μg psPAX2 (Addgene #12260) in 1.5 mL Opti-MEM (Gibco) and transfected using 60 μL TransIT-293 reagent (Mirus Bio, 2700). After a 30-min incubation at room temperature, the transfection mixture was added dropwise to cells at 80–90% confluency in ESC medium without LIF. Cells were incubated for 48 h at 37 °C and 5% CO₂. Lentiviral supernatant was collected and clarified either through a 0.45-μm PES filter (Sartorius, 16533) or by centrifugation at 500 × *g* for 10 min. Virus was concentrated using Lenti-X Concentrator (Takara, 631231) at a 1:3 volume ratio, incubated at 4 °C for 4–16 h, and centrifuged at 1,500 × *g* for 45 min at 4 °C. Viral pellets were resuspended in ESC medium at 1/10–1/20 of the original supernatant volume and stored at −70 °C.

### Lentiviral titration

CRISPRi ESCs expressing dCas9-KRAB-MeCP2 (Addgene #110821) were maintained under blasticidin (InvivoGen, ant-bl-1) and hygromycin (InvivoGen, ant-hg-1) selection for 2–3 passages before lentiviral titration. Cells (1.2 × 10^5^ per well) were transduced with 0–20 μL of sgRNA lentivirus in ESC medium containing 8 μg/mL polybrene. Non-transduced controls with and without subsequent puromycin (InvivoGen, ant-pr-1) selection were included. At 48 h post-transduction, cells were passaged and expanded. The following day, a fraction of each condition was analyzed by flow cytometry following DAPI staining, with transduced cells identified as DAPI−/mCherry+. The mCherry+ gate was defined using the non-transduced control, and the proportion of mCherry+ cells was used to estimate multiplicity of infection (MOI). Parallel cultures were selected with 1 μg/mL puromycin for 3 days, after which the non-transduced control contained no viable cells and transduced cultures were 98–100% mCherry+. A lentiviral input of 10 μL per 1.2 × 10^5^ cells (∼120 μL per 10^6^ cells) produced an MOI of approximately 10% and was used for subsequent CRISPRi experiments.

### Immunofluorescence

Cells were trypsinised, washed with PBS, and fixed in 2% paraformaldehyde for 30 min at room temperature. Fixed cells (100 μL) were cytospun onto Polysine slides for 3 min at 200 × *g*, permeabilised with 0.5% Triton X-100 for 30–60 min, and blocked in PBS containing 0.05% Tween-20 and 1% BSA for 30–60 min at room temperature or overnight at 4 °C. Slides were incubated for 1 h at room temperature with rabbit anti-SOX1 (1:500; Abcam, ab227118) or goat anti-SOX2 (1:100; R&D Systems, AF2018), followed by anti-rabbit Alexa Fluor 568 or anti-goat Alexa Fluor 488 secondary antibody (1:500), respectively, for 1 h in the dark. Slides were stained with 5 μg/mL DAPI for 15 min, then mounted using SlowFade Gold mounting medium (Molecular Probes). Images were acquired on a Zeiss LSM 780 confocal microscope using 40× or 63× oil-immersion objectives and processed using Zeiss ZEN or FIJI software.

### RNA isolation and qRT-PCR

Cells were washed in PBS then lysed in 350–600 μL buffer RLT (Qiagen) to release the RNA. Total RNA was isolated using the RNeasy Mini Kit (Qiagen) and 0.1 – 1 μg TURBO DNAse-treated (Thermo Fisher AM1907) RNA was converted to first strand cDNA using random hexamer priming (Thermo RevertAid K1622). Quantitative reverse-transcription PCR (qRT-PCR) was performed according to manufacturer’s instructions using Brilliant III SYBR master mix (Agilent Technologies), and relative quantification was performed using the comparative cycle threshold (CT) method with normalization to *CycloB1* levels. Primer sequences for validated qRT-PCR primers are presented in **Supplementary Table 1**. Primers were generated for this study using Primer-BLAST (ncbi.nlm.nih.gov/tools/primer-blast/) and synthesised by Sigma-Aldrich.

### Western blotting

Protein lysates were generated by resuspending cell pellets in Laemmli buffer (62.5 mM Tris-HCl pH 6.8, 2% SDS, 5% β-mercaptoethanol, 10% glycerol) and incubating for 3 – 5 min at 98 °C. Protein concentration was determined by Bradford Assay (Bio-Rad). Boiled samples were loaded onto NuPAGE 4 – 12% Bis-Tris gels (Invitrogen NP0322BOX) in NuPAGE MES SDS (Invitrogen NP0002) or MOPS SDS running buffer (Invitrogen NP0001), then blotted on polyvinyl difluoride membranes. Blocking was performed using 5% skimmed milk/0.01%Tween/PBS, then membranes were incubated with primary antibodies for 16 h (anti-β-ACTIN Abcam ab6276 1:20,000; anti-FLAG M2 Sigma-Aldrich F1804 1:1000; anti-H2A.X Abcam ab200669 1:1000; anti-HSP90 CST 4874 1:1000; antimacroH2A.2 (H2AFY2) Proteintech 17030-1-AP 1:1000; anti-pH2A.X (Ser139) Sigma-Aldrich clone JBW301 1:1000; anti-TET1 [N1] GeneTex GTX125888 1:1000). Samples were washed in 0.01%Tween/PBS (3× briefly, 3× > 10 min), incubated >1 h with horseradish peroxidase-conjugated secondary antibodies (anti-mouse, Santa Cruz sc-2005, 1:5000; anti-rabbit, Bio-Rad 170–6515, 1:5000), then washes were repeated. Detection was performed with enhanced chemiluminescence reagent (GE Healthcare, RPN2209).

### Co-immunoprecipitation

Nuclear extracts were prepared using buffer A (10 mM HEPES, 1.5 mM MgCl₂, 10 mM KCl, 0.5 mM DTT, 0.05% NP-40, pH 7.9) and buffer B (5 mM HEPES, 1.5 mM MgCl₂, 0.2 mM EDTA, 0.5 mM DTT, 26% glycerol, pH 7.9), supplemented immediately before use with 250 U Benzonase and complete protease inhibitors (Roche). Cells from 15-cm dishes were scraped into 500 μL ice-cold buffer A, incubated on ice for 10 min, and centrifuged at 3000 rpm for 10 min at 4 °C. Nuclear pellets were resuspended in 374 μL buffer B, supplemented with 26 μL 4.6 M NaCl, homogenized with Dounce or glass homogeniser, incubated on ice for 30 min, and centrifuged at 24,000 × *g* for 20 min at 4 °C. Protein concentrations of the resulting nuclear extracts were determined by Bradford assay (Bio-Rad).

For immunoprecipitation, 50 μL Protein G Dynabeads (Invitrogen) were washed and incubated with 5 μg antibody in 500 μL wash buffer (10 mM Tris-HCl, pH 7.5, 150 mM NaCl, 0.5 mM EDTA) for 1–4 h at 4 °C. Antibody-bound beads were washed three times and incubated with 250 μg nuclear protein in a final volume of 500 μL for 1–16 h at 4 °C with end-over-end rotation; 10% of each lysate was retained as input. Beads were washed three times and proteins were eluted in 50 μL Laemmli buffer by heating for 10 min at 75 °C. Immunoprecipitated proteins (20 μL eluate) and nuclear lysates (10–25 μg) were analyzed by western blotting.

### RIME and qPLEX-RIME

Cell pellets (140 mg per replicate) were fixed, lysed, and immunoprecipitated as described for ChIP using 5 μg anti-FLAG M2 antibody (Sigma-Aldrich, F1804). Following immunoprecipitation, beads were washed ten times with RIPA buffer and twice with 100 mM ammonium bicarbonate. Samples were processed by the CRUK Cambridge Institute Proteomics Core for RIME and qPLEX-RIME as previously described (55). Briefly, proteins were digested on-bead with trypsin, peptides were C18-cleaned and labelled with TMT 10-plex reagents (Thermo Scientific), and samples were pooled and fractionated. Peptides were analyzed using a Dionex Ultimate 3000 UHPLC system coupled to a Fusion Lumos mass spectrometer (Thermo Fisher Scientific).

### Full Proteome

The cell pellets were resuspended in the lysis buffer containing 100 mM triethylammonium bicarbonate (TEAB, Sigma), 10% isopropanol, 50 mM NaCl, 1% SDC with nuclease and protease/phosphatase Inhibitors, followed by 15 min incubation at room temperature (RT). Total proteins were reduced and alkylated with 2 μL of 50mM tris-2-caraboxymethyl phosphine (TCEP, Sigma) and 1 μl of 200 mM iodoacetamide (IAA, Sigma), respectively, for 1 hour. Then protein samples were digested overnight at 37 °C in a trypsin solution at a protein/trypsin ratio of ∼1:30. The next day, protein digest was labelled with the TMT pro reagents (Thermo Scientific) for 1 hour. Then protein samples were digested overnight at 37 °C in a trypsin solution at a protein/trypsin ratio of ∼1:30. The next day, protein digest was labelled with the TMT pro reagents (Thermo Scientific) for 1 hour and quenched with 5% hydroxylamine (Thermo Scientific) for 15 min at RT. All the samples were mixed and acidified using formic acid for SDC removal. The dry TMT mix was fractionated on a Dionex Ultimate 3000 system at high pH using the X-Bridge C18 column (3.5 μm, 2.1x150mm, Waters) with 90 min linear gradient from 5% to 95% acetonitrile contained 20 mM ammonium hydroxide at a flow rate of 0.2 ml/min. Global proteome fractions were reconstituted in 0.1% formic acid for liquid chromatography tandem mass spectrometry (LC–MS/MS) analysis.

### qPLEX-RIME data processing

qPLEX-RIME data were processed by the CRUK Cambridge Institute Proteomics Core as described previously (55). Spectra were searched against the UniProt *Mus musculus* proteome (taxon ID 10090), with tandem mass tag modifications specified as appropriate. Peptide identifications were filtered at a 1% false discovery rate (FDR), and uniquely mapping peptides were used for protein quantification. Data normalization and statistical analysis were performed using the Bioconductor package qPLEXanalyzer (55). Peptide intensities were median-normalized and summed to obtain protein-level abundances. Differential protein abundance was assessed using limma (56), with Benjamini–Hochberg correction for multiple testing (57).

### RNA-seq

RNA was isolated as described above. Opposite strand-specific polyA RNA-seq libraries were prepared from 1 μg DNase-treated RNA with RNA integrity number (RIN) ≥8 using the Sanger Institute Illumina bespoke pipeline and sequenced as 150-bp paired-end reads on an Illumina HiSeq 2000. Reads were trimmed with Trim Galore (v0.6.6) and aligned to the *Mus musculus* GRCm38 genome using HISAT2. BAM files were imported into SeqMonk using standard parameters, and strand-specific mRNA abundance was quantified using the SeqMonk RNA-seq quantification pipeline. Quantified data were exported to R for downstream analysis.

### ChIP-seq

Cells were fixed in 1% formaldehyde (Fisher Scientific 28906) and 2 mM DSG (Thermo Scientific A35392) in DMEM (Invitrogen 41966-052) or PBS for 10 min, then glycine was added to 0.125 M to quench. Cells were scraped, normalized to 80 mg of cell pellet per replicate, then lysed with buffer LB1 (50 mM HEPES pH 7.5, 140 mM NaCl, 1 mM EDTA, 10% Glycerol, 0.5% Igepal CA-630, 0.25% Triton-X 100) on ice for 10 min, LB2 (10 mM Tris-HCl pH 8.0, 200 mM NaCl, 1 mM EDTA, 0.5 mM EGTA) on ice for 5 min, then resuspended in LB3 (10 mM Tris-HCl pH 8.0, 100 mM NaCl, 1 mM EDTA, 0.5 mM EGTA, 0.1% Na-Deoxycholate, 0.5% N-lauroylsarcosine, 1× complete protease inhibitors). Chromatin was fragmented by sonication for 25–35 cycles of 30 s ON high/30 s OFF (Diagenode Bioruptor). For each ChIP, 5 μg anti-FLAG M2 Sigma-Aldrich F1804 (1:1000; Sigma-Aldrich F1804) was prebound to protein G dynabeads (Invitrogen), then blocked with 0.5% BSA in PBS. Sonicated DNA was added to antibody-bound beads with 10% Triton-X 100, then incubated and rotated at 4 °C for 16 h. Samples were washed 6 times with RIPA buffer (50 mM HEPES pH 7.6, 1 mM EDTA, 0.7% Na-deoxycholate, 1% Igepal CA-630, 0.5 M LiCl), then washed once in 1× TBS, then incubated at 65 °C in elution buffer (50 mM Tris-HCl, pH 8; 10 mM EDTA; 1% SDS) for 16 h. Samples were treated with RNase A and Proteinase K then DNA purification was performed with MinElute PCR purification columns (Qiagen 28006). Following ChIP and DNA purification, library quality was assessed using Agilent Bioanalyzer High Sensitivity DNA chips. ChIP-seq libraries were prepared using the CRUK Cambridge Institute Illumina bespoke pipeline. Libraries were sequenced as 50-bp paired-end reads on an Illumina NovaSeq X. Reads were trimmed with Trim Galore (v0.6.6) and aligned to the *Mus musculus* GRCm38 genome using Bowtie2.

### ChIP-seq peak calling

ChIP-seq peaks were called independently for each replicate using MACS2 callpeak with default parameters; the --broad option was used for macroH2A.2, H1c, and H1d.

Unless otherwise stated, promoters were defined as −2 kb to +0.5 kb relative to the transcription start site. TET1 ChIP-seq data were generated and processed as described above. Published raw ChIP-seq datasets for H1c, H1d, macroH2A.2, pH2A.X, SIN3A, SSRP1, SUPT16, and SUZ12 were reprocessed using the same alignment and peak-calling workflow (44, 45, 58–62).

### Comparative RNA-seq analysis of *in vivo* and i*n vitro* neuroectoderm

Processed single-cell RNA-seq data from *in vivo* mouse gastrulation were obtained using the MouseGastrulationData R package (63). Cells were restricted to neuroectoderm-associated populations (Epiblast, Rostral neurectoderm, Caudal epiblast, Caudal neurectoderm, Forebrain/Midbrain/Hindbrain, Spinal cord, NMP, and Neural crest), and pseudobulk profiles were generated by aggregating cells of each cell type within each sample. Bulk RNA-seq data from *in vitro* neuroectoderm differentiation were processed as described above and restricted to biological replicates from Days 1–6 (51). Differential expression analysis was performed separately for each dataset using DESeq2, with Benjamini–Hochberg-adjusted *P* < 0.05 considered significant. Variance-stabilizing transformation was used for visualization. Gene sets derived from the top 250 and 1,000 most variable genes in the *in vivo* and *in vitro* datasets, respectively, were compared to identify shared and dataset-specific genes. Gene Ontology enrichment analysis was performed using PANTHER and GOliath with an FDR threshold of 5%.

### Knockdown and Knockout RNA-seq analysis

Published knockdown and knockout RNA-seq datasets were processed and quantified as described above (31, 60, 62). Differential expression between wild-type and perturbed samples was assessed using DESeq2, with log_2_ fold-change shrinkage performed using apeglm (64, 65).

### Chromatin state analysis

Published ENCODE peak sets for CTCF, DNase hypersensitivity, H3K4me1, H3K4me3, H3K9ac, H3K9me3, H3K27ac, H3K27me3, H3K36me3, NANOG, and POU5F1 were used to generate genome-wide chromatin-state annotations using a 10-state ChromHMM model with default parameters (66, 67). Factor peaks were assigned to overlapping chromatin states. For promoters overlapping multiple states, the active, bivalent, primed, or repressed state overlapping or nearest the transcription start site (TSS) was assigned. Chromatin-state enrichment was calculated relative to the genomic frequency of each state.

### ChIP-seq analysis

ChIP-seq analyses were performed in R. Peaks were annotated to the mm10 genome using annotables, TxDb.Mmusculus.UCSC.mm10.knownGene, and ChIPseeker (68). Canonical transcript TSSs were identified using the transcript_is_canonical attribute in biomaRt (69). Genomic overlaps were calculated using subsetByOverlaps from GenomicRanges (70). Venn diagrams were visualized using venneuler, and heatmaps of ChromHMM-state overlaps were generated using ComplexHeatmap (71).

### Integrated ChIP-seq and RNA-seq analysis

Gene Ontology enrichment analysis was performed using the biological process ontology in clusterProfiler with otherwise default parameters (72). Heatmaps were generated using ComplexHeatmap (71), and additional plots were made using ggplot2, ggrepel, and ggsignif (73). Statistical analyses were performed in R using the tests and parameters specified in the main text.

## RESULTS

### Proteome and TET1 chromatin complex dynamics during the transition from pluripotency to early NPCs

We performed multiomic profiling of TET1 chromatin complex dynamics during an *in vitro* NPC differentiation timecourse, including whole-cell proteomics, qPLEX-RIME to measure TET1-associated chromatin proteins, and ChIP-seq to measure TET1 chromatin occupancy (**Figure 1A**), using mESCs expressing endogenous TET1 with a C-terminal 3×FLAG tag (26, 55, 74). The NPC differentiation model recapitulated gene-expression patterns observed during *in vivo* neuroectoderm development (**Supplementary Figure 1**). Proteome and qPLEX-RIME measurements were reproducible across replicates and showed marked differentiation-dependent changes (**Figure 1B–F; Supplementary Figures 2–3**). Between Day 0 and Day 2, 3,724 proteins were differentially expressed (adjusted p < 0.05; **Figure 1D**). Neuroectoderm-priming factors PRTG and SOX3 were among the 20 most upregulated proteins, whereas the pluripotency regulators JARID2, POU5F1, and TCEA3 decreased by more than 1.5-fold (75–79). Together with the RNA-seq time course (**Supplementary Figure 1**), these changes establish Day 2 as an early NPC state.

**Figure 1.**
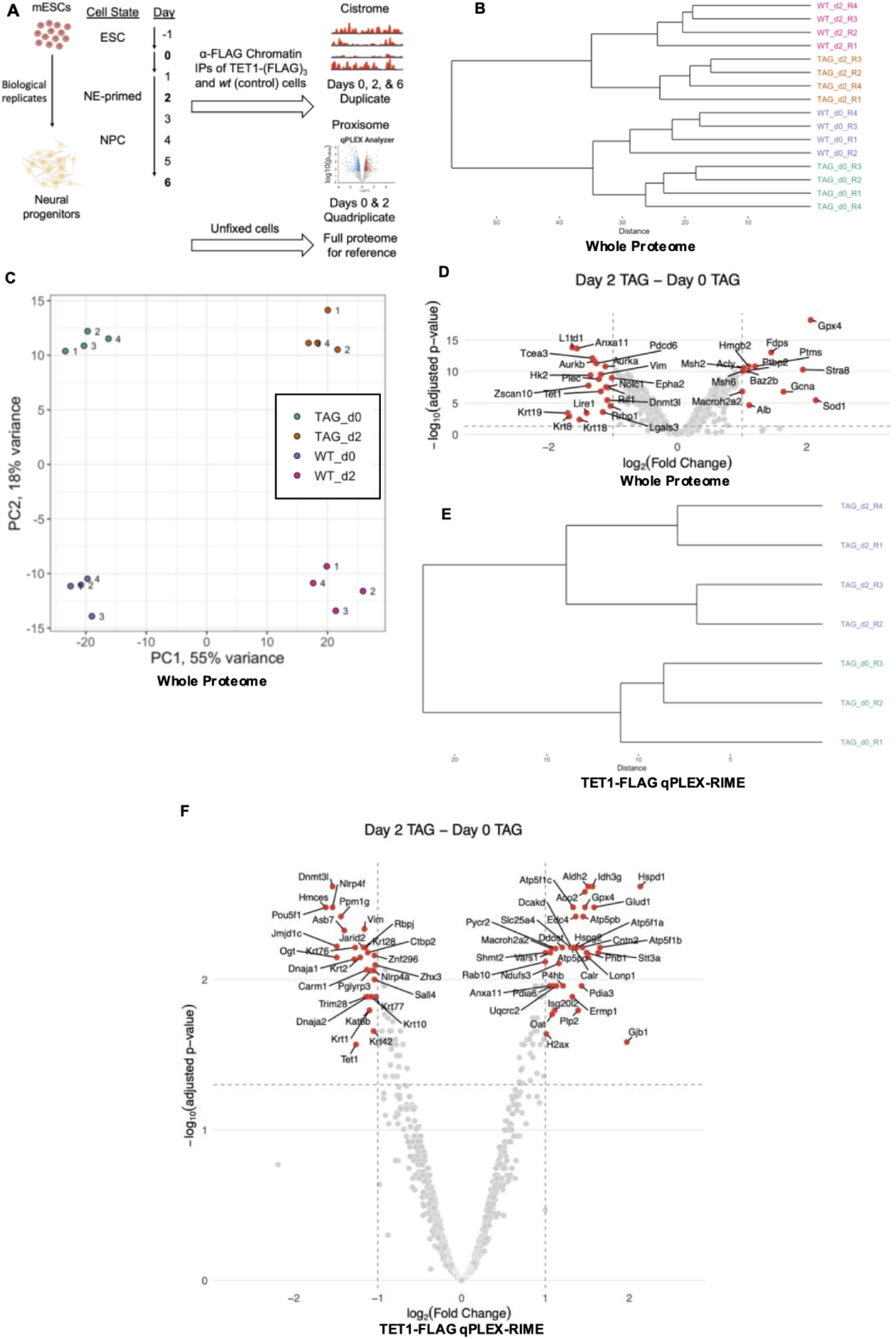
Proteome and TET1 proxisome dynamics during NPC differentiation. **A.** Experimental scheme for determining TET1 functional interactions on chromatin during *in vitro* NPC differentiation by qPLEX-RIME, ChIP-seq, and RNA-seq. **B.** Hierarchical clustering of normalized protein intensities from proteome samples using euclidean distance and complete linkage methods. **C.** PCA of normalized protein intensities from proteome samples. **D.** Volcano plot of differentially expressed proteins between Day 2 TAG versus Day 0 TAG proteome (quadruplicate). A limma-based statistical analysis was conducted. Differentially expressed proteins (adjusted p-value < 0.05, |LFC| > 1.5) are coloured red and labelled. **E.** Clustering of normalized protein intensities using euclidean distance and complete linkage methods after batch effect correction using Combat. **F.** Volcano plot of differentially enriched proteins between Day 2 TAG versus Day 0 TAG qPLEX-RIME replicate samples after removal of proteins not significantly enriched compared to WT control. A limma-based statistical analysis was conducted. Differentially enriched proteins (adjusted p-value < 0.05, |LFC| > 2) are coloured red and labelled. ESC = embryonic stem cell. NE = neuroectoderm. NPC = neural progenitor cell. TAG = TET1-(FLAG)_3_ cell line. WT = wild-type cell line. LFC = log fold-change.

Between mESCs and early NPCs, 128 proteins were differentially enriched by qPLEX-RIME (**Figure 1F**). POU5F1 and JARID2, which co-occupy chromatin with TET1 in pluripotent cells, were among the most strongly depleted proteins (30, 45). Thus, the TET1 proxisome reorganised as cells transitioned from pluripotency to an early NPC state. TET1 protein abundance decreased approximately twofold upon pluripotency exit, potentially reducing the detection of qPLEX-RIME interactions in early NPCs (**Figures 1F, 2A**). We therefore focused our analysis on proteins enriched more than 1.5-fold by qPLEX-RIME in early NPCs and differentially expressed in the whole-cell proteome, while recognizing that reduced bait abundance could produce false negatives. We further required evidence of nuclear localization (80), expression during *in vivo* neuroectoderm differentiation (63), and chromatin binding (**Supplementary Figure 4A–F**). This qPLEX-RIME data filtering resulted in five candidates: H2A.X, macroH2A.2, PTMS, PHB1, and EDC4 (81–84). We next tested candidates for physical association with TET1 and a role in NPC differentiation.

**Figure 2.**
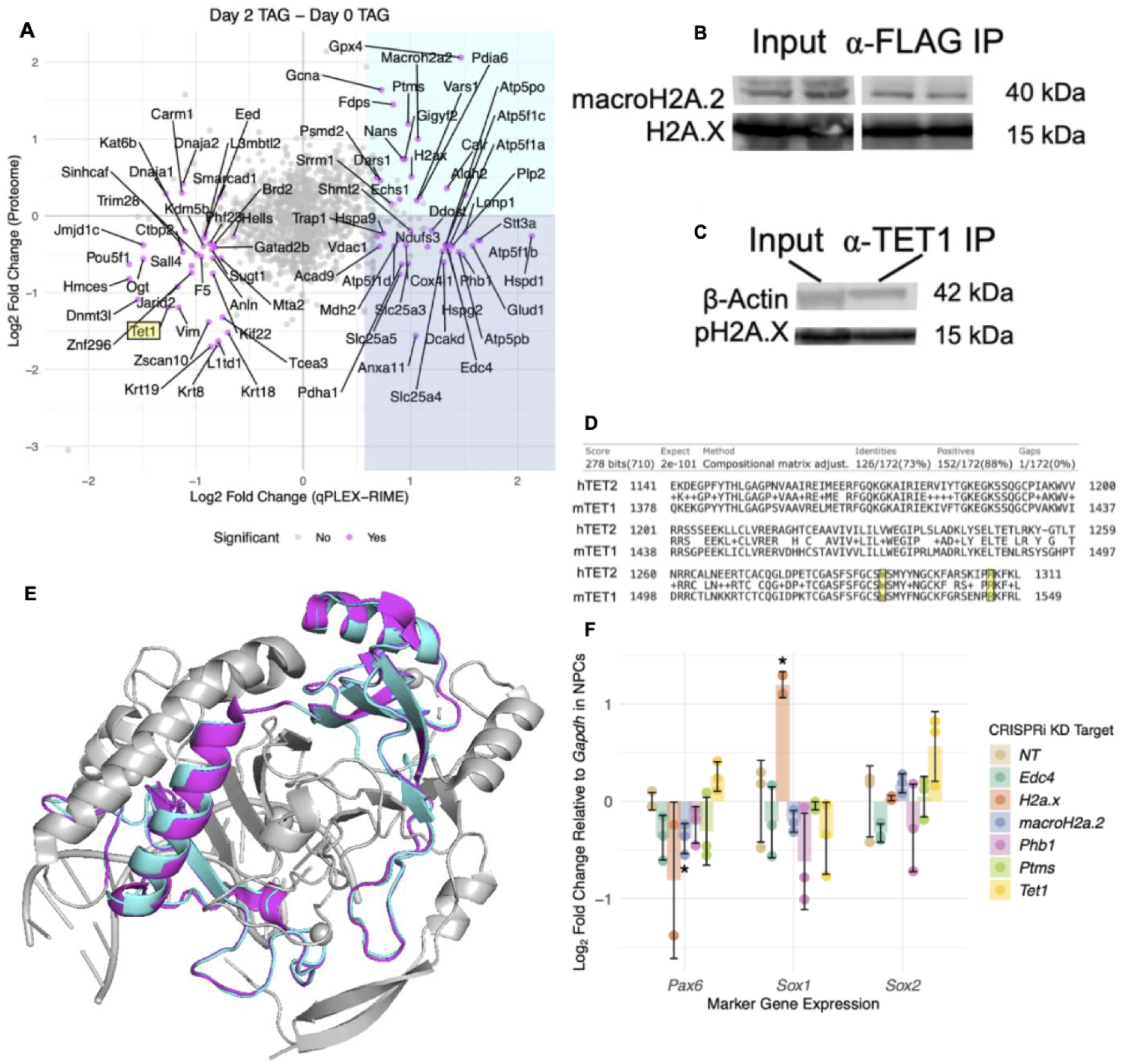
TET1 interacts with H2A.X and macroH2A.2 during NPC differentiation. **A.** Changes in protein expression (y-axis; full proteome) versus changes in TET1-associated chromatin complex proteins (x-axis; qPLEX-RIME) between Day 2 versus Day 0 of NPC differentiation. All differentially expressed (adjusted p-value < 0.05) and enriched proteins are highlighted magenta. The upregulated (LFC>0) and enriched (LFC>1.5) region is highlighted cyan (16 proteins). Downregulated (LFC<0) and enriched (LFC>1.5) region is highlighted blue (28 proteins). TET1 is highlighted yellow. **B–C.** TET1 directly interacts with H2A.X, pH2A.X and macroH2A.2 in mESCs. Representative co-IPs of TET1-(FLAG)_3_ mESCs involving either an α-FLAG (**G**) or α-TET1 IP (**H**) followed by blots for H2A variants**. D.** pBLAST sequence alignment of mTET1 (UniProt Accession: Q3URK3) Cys-rich domain (aa 1378–1549) to hTET2 (UniProt Accession: Q6N021) Cys-rich domain (aa 1141–1549) demonstrates high homology. The conserved tryptophan (W) residue required for hTET2 H3K36me1/2 specificity and arginine (R) residue required for (M)CGW DNA substrate specificity are highlighted. **E.** Alignment of TET1 Cys-rich domain structure (cyan; AlphaFold ID: AF-Q3URK3-F1-v4) to the experimentally determined hTET2 structure bound to DNA (pink; PDB ID: 4NM6) demonstrates that the domain’s structure is conserved. The rest of the TET2 structure bound to DNA is coloured gray. **F.** Effects of CRISPRi KD on NPC differentiation of candidate functional interactors with TET1. rt-qPCR of NPC marker genes demonstrates CRISPRi-mediated knockdown of *H2a.x* and *macroH2a.2* promote and impair NPC differentiation, respectively. Experiments performed in triplicate (duplicate for *H2a.x* due to one replicate failing QC). WT NPCs as negative control. Error bars indicate standard error. A two-sided t-test was performed. An asterisk denotes a p- value < 0.05. LFC = log fold-change.

### MacroH2A.2 and pH2A.X physically and functionally interact with TET1 during NPC differentiation

MacroH2A.2, H2A.X, and Ser139-phosphorylated H2A.X (pH2A.X), which is a post-translational modification that promotes DNA double strand break repair and pluripotency self-renewal (83, 85–88), were detected in native TET1 immunoprecipitates from mESCs (**Figure 2B-C**). The other qPLEX-RIME candidates were not detected by co-IP. We searched for possible interaction domains between TET1 and H2A variants. The DNA-binding Domain (DBD) in the Cys-rich (CR) domain of the globular catalytic core of human TET2 binds H2A, H2B, and H3, and recognises H3K36me1/2 (Yamagata and Kobayashi, 2017). The mouse TET1 CR domain shares 73% sequence identity and 88% sequence similarity with the homologous human TET2 domain, adopts a nearly identical AlphaFold-predicted structure (89, 90), and conserves the region surrounding W1291 involved in H3K36me binding (**Figure 2D-E**). R1302, which contributes to human TET2 DNA-sequence specificity, is also conserved (**Figure 2D**), consistent with shared sequence preferences among mouse and human TET enzymes (91). Together, these biochemical, sequence and structural data indicate that the TET1 CR domain directly interacts with histones and may couple H3K36me1/2 recognition to chromatin localization.

We used CRISPRi-mediated silencing followed by qRT-PCR to test whether qPLEX-RIME candidates regulated NPC differentiation. *Tet1* knockdown was incomplete for all gRNAs tested and resulted in no significant changes in NPC-marker gene expression after directed NPC differentiation (**Figure 2F**; **Supplementary Figure 4G**). In contrast, strong CRISPRi-mediated *macroH2A.2* knockdown reduced *Pax6* expression, whereas *H2A.X* knockdown increased *Sox1* expression (**Figure 2F**). These results agree with prior findings that macroH2A.2 promotes NPC differentiation whereas H2A.X limits neuroectoderm differentiation potential (86, 92, 93).

### TET1 and H2A variants co-occupy distinct chromatin states in pluripotency

To define regions of TET1 and H2A co-occupancy in mESCs, we performed TET1 ChIP-seq and compared our data to published H2A variant ChIP-seq. MACS2 called 1042 and 22313 peaks for published pH2A.X and macroH2A.2 ChIP-seq datasets, respectively, whereas available H2A.X ChIP-seq datasets had insufficient enrichment or replication for peak calling (59, 61, 94, 95). In mESCs, 13.3% of pH2A.X peaks and 17.1% of macroH2A.2 peaks overlapped TET1 peaks (**Figure 3A-D**). In contrast, only 2.0% of pH2A.X peaks overlapped macroH2A.2, indicating that the two variants rarely co-occupy chromatin.

**Figure 3.**
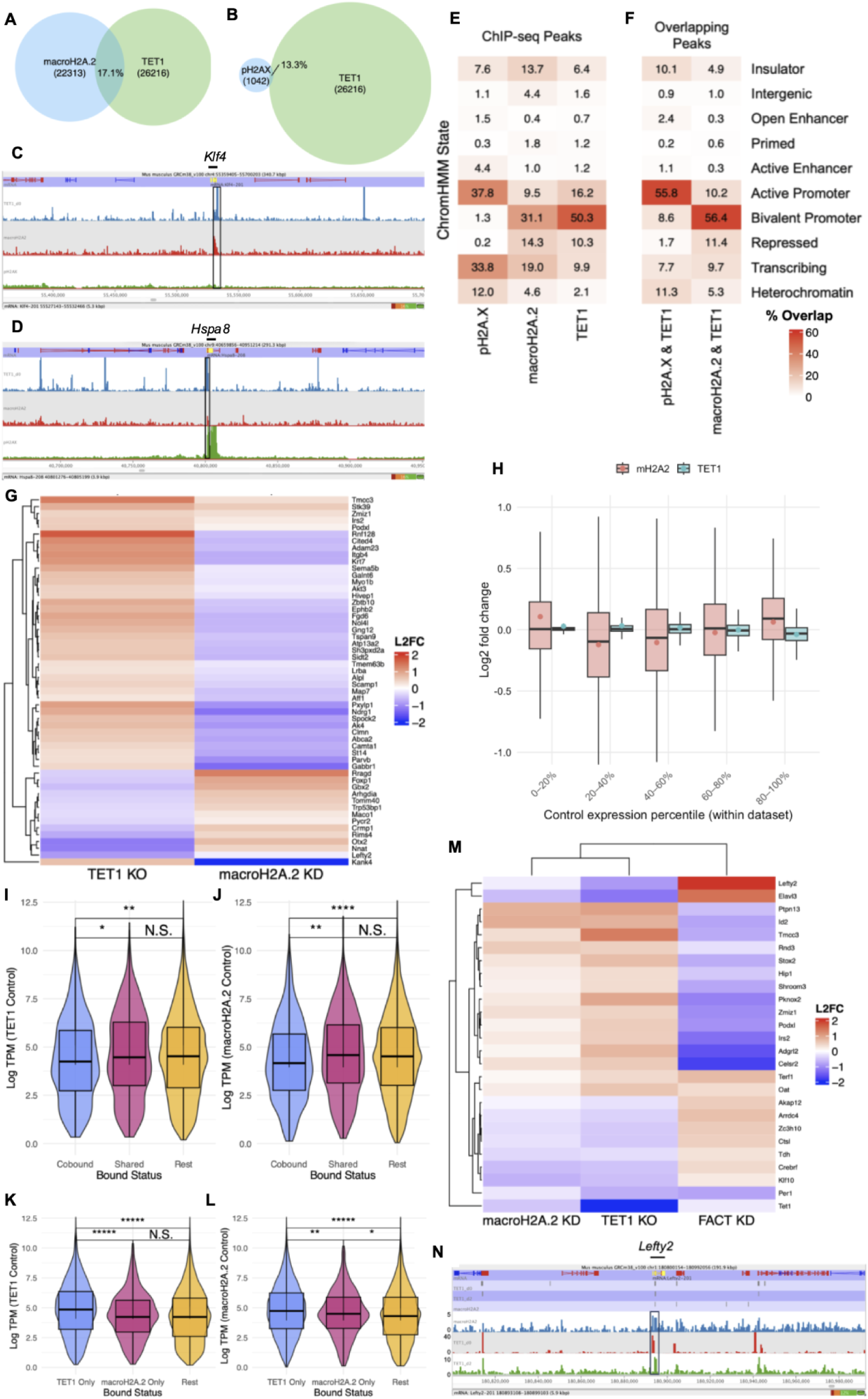
TET1 and macroH2A.2 functional antagonism in mESCs. **A.** Venn Diagrams of ChIP-seq peak overlap between macroH2A.2 (**A**) or pH2A.X (**B**) and TET1. **C–D.** Genome browser tracks of macroH2A.2 (**C**) and pH2A.X (**D**) ChIP-seq peak overlap with TET1 ChIP-seq Peaks in mESCs. **E.** ChIP-seq Peaks and **F.** Overlapping ChIP-seq Peaks at genomic regulatory elements in ESCs for pH2A.X, macroH2A.2, and TET1 shows pH2A.X and TET1 enrich in transcriptionally active elements while macroH2A.2-TET1 enrich in active and bivalent promoters and Polycomb-repressed domains. Overlap with ChromHMM states is normalised such that each ChIP-seq Peak set sums to 100% across ChromHMM states. **G.** Log2 FC expression of shared high-confidence DEGs (FDR < 0.1, |Log FC| > 1.5) in TET1 KO and macroH2A.2 KO versus wt mESCs (van der Veer et al., 2023) at TET1 and macroH2A.2 ChIP-seq overlapping targets demonstrates a general pattern of opposite effects by TET1 and macroH2A.2 on gene expression. Genes are clustered by expression pattern. **H.** Boxplots of log2 fold change in gene expression separated into log TPM quintile demonstrates that TET1 becomes activating at highly expressed genes while macroH2A.2 switches from activating to repressive as gene expression increases across the upper 4 quintiles. The mean is represented by a point while the median is represented by a solid black line within each boxplot. **I.** Gene-level log TPM for co-bound versus shared versus the rest of gene promoters in WT mESCs (van der Veer et al., 2023 dataset). **J.** Gene-level log TPM for co-bound versus shared versus the rest of gene promoters in WT mESCs (Valakos et al., 2023 dataset). Cobound = TET1 and macroH2A.2 ChIP-seq peaks overlap. Shared = TET1 and macroH2A.2 ChIP-seq peaks are both present but do not overlap. Rest = all other promoters. A promoter is defined as the region from 2kb upstream to 0.5 kb downstream of a TSS. **K.** Gene-level log TPM values at TET1-only promoters, macroH2A.2-only promoters, versus and the rest of gene promoters in WT mESCs (van der Veer et al., 2023 dataset). **L.** Gene-level log TPM values at TET1-only promoters, macroH2A.2-only promoters, and the rest of gene promoters in WTmESCs (Valakos et al., 2023 dataset). For panels **I–L**, a Kruskal-Wallis rank sum test was performed to establish significance followed by a post-hoc Dunn’s test of multiple comparisons. Asterisk indicates adjusted p-value < 0.05; each additional asterisk indicates an order of magnitude decrease in p-value. **M.** Log2 FC expression of shared DEGs (FDR < 0.1) in TET1 KO, macroH2A.2 KD, and FACT KD versus WT mESCs in which TET1 KO and macroH2A.2 KD cause same-direction gene expression changes demonstrates a pattern of FACT functantagonism. Genes are clustered by expression pattern. **N.** TET1 and macroH2A.2 binding of the *Lefty2* promoter. Genome browser (Seqmonk) view displaying TET1 ChIP-seq tracks from Day 0 and Day 2 of NPC differentiation, along with a macroH2A.2 ChIP-seq track from Day 0.

We determined the chromatin contexts of TET1–H2A-variant co-occupancy by intersecting TET1, pH2A.X, and macroH2A.2 ChIP-seq peaks with ChromHMM states in mESCs (**Supplementary Figure 5A–B; Figure 3C–D**). TET1 was most strongly associated with bivalent promoters (50.3%), followed by active promoters (16.2%; **Figure 3E**). More than 70% of pH2A.X enrichment occurred in active promoters (37.8%) or transcribed chromatin (33.8%), whereas macroH2A.2 was most strongly associated with bivalent promoters (31.1%) but was also represented in transcribed (19%), repressed (14.3%), and active regions (9.5%; **Figure 3E**). Co-occupancy with TET1 occurred in two distinct promoter contexts: TET1–pH2A.X co-occupied active promoters and TET1–macroH2A.2 co-occupied bivalent promoters (**Figure 3F**). The TET1-dependent corepressors SIN3A and SUZ12 are also enriched at TET1-bound bivalent promoters (**Supplementary Figure 5C–D**). Together with their physical association (**Figure 2B–C**), these findings are consistent with an interaction between TET1 repressive protein complexes and macroH2A.2 in bivalent chromatin (29). In contrast, TET1 and pH2A.X associate within transcriptionally active chromatin through a shared upstream mechanism (96, 97).

### TET1 and macroH2A.2 exert opposing chromatin context-dependent effects in pluripotency

To determine how the chromatin contexts of these factors related to their transcriptional regulation, we integrated published TET1 KO and macroH2A.2 KD RNA-seq with the ChIP-seq and ChromHMM datasets (31, 62). Among high-confidence differentially expressed genes (DEGs; absolute fold change > 1.5; FDR < 0.1), TET1 predominantly functioned as a repressor in repressive chromatin contexts, consistent with previous reports (29), whereas macroH2A.2 tended to increase expression, especially in primed chromatin (**Supplementary Figure 5E–F**). Among high-confidence DEGs with promoters co-bound by TET1 and macroH2A.2, TET1 repressed 76% (41/54) of DEGs, and 89% (48/54) of DEGs responded in opposite directions upon TET1 or macroH2A.2 depletion (**Figure 3G**). This pattern extended to the entire set of overlapping DEGs, consistent with global regulatory effects of these factors producing a functional rather than direct antagonism; indeed, both factors activated a small shared group of active loci and silenced a small shared group of active and bivalent loci (**Supplementary Figure 5G–H**). Across expression strata, TET1 became progressively more activating as basal expression increased, consistent with its established activator role at highly expressed housekeeping genes (29), whereas macroH2A.2 showed the opposite trend across the upper four quintiles (**Figure 3H**). Co-occupancy was associated with lower expression compared to separate occupancy (**Figure 3I–L**). Together, these results demonstrate that TET1 and macroH2A.2 act as opposing, context-dependent transcriptional rheostats that co-occupy bivalent promoters and lowly expressed genes.

### MacroH2A.2 and H1 occupy mutually exclusive chromatin regions

One potential mechanism for macroH2A.2 to activate lowly expressed genes is by displacement of linker histone H1, as previously demonstrated in chick chromatin (98). The macroH2A linker can compete with H1 for the nucleosomal DNA entry–exit site, but the macro domain prevents full-length macroH2A from reproducing H1’s stabilizing effect on nucleosomes (99–103). Consistent with this model, we found that ChIP-seq peaks of H1c and H1d—the predominant H1 isotypes in mESCs—rarely overlapped macroH2A.2 peaks in non-intergenic chromatin and did not overlap at bivalent regions: only 1.0% of H1c peaks and 0.5% of H1d peaks overlapped macroH2A.2 within annotated non-intergenic states (58, 104–106). MacroH2A.2 may therefore support the exclusion of H1 from lowly expressed developmental promoters, reducing nucleosome stabilization and thus poising them for activation.

### FACT opposes TET1–macroH2A.2 concordant regulation

We tested for an association between H2A exchange and TET1–macroH2A.2 regulation by integrating published FACT ChIP-seq and FACT KD RNA-seq with our TET1 KO and macroH2A.2 KD analyses (60). FACT is an H2A/H2B chaperone involved in macroH2A.2 removal and H2A.X exchange (83, 107, 108). Its core subunits, SSRP1 and SUPT16H, were enriched in TET1 qPLEX-RIME data in mESCs and early NPCs, and 31.0% of FACT-bound regions overlapped TET1 peaks (**Supplementary Figure 3K–L**). These results place FACT, TET1 and H2A variants in a shared chromatin-regulatory network.

Although TET1 and macroH2A.2 regulated most co-bound genes in opposite directions, their depletion produced concordant up- or downregulation in 25 DEGs co-regulated by FACT. FACT depletion caused opposite changes in 88% (22/25) of these DEGs (**Figure 3M**). These 22 DEGs included the developmental regulators *Id2* and *Shroom3*, which were repressed by TET1 and macroH2A.2, and the Nodal antagonist *Lefty2* (**Figure 3N**), which was activated by both factors and is known to act downstream of TET1 catalysis to inhibit primitive streak formation (22, 24, 28, 30, 31, 109, 110). In line with these findings, factor depletion experiments demonstrate that FACT opposes NPC differentiation from pluripotency while TET1 and macroH2A.2 promote it (31, 60, 92).

### TET1 represses alternative regulatory programs in early NPCs

We performed TET1 ChIP-seq across NPC differentiation and called 26,216 peaks in mESCs and 6,581 peaks in early NPCs. No peaks passed the MACS2 calling threshold at our final timepoint, Day 6, consistent with an early role for TET1 in promoting NPC differentiation before it is sharply downregulated (30, 31, 106). In early NPCs, when TET1 RNA abundance was approximately one quarter of its level in pluripotent cells, 81% (21150/26216) of peaks had been lost and relatively few (20%, 1347/6581) new sites had appeared (**Supplementary Figure 6A–E**). Despite this global loss, the proportion of TET1 peaks near transcription start sites increased from 48% in mESCs to 75% in early NPCs, and 80% of sites retained across both timepoints were within promoters (**Supplementary Figure 6C-E**). TET1 was also lost from poised enhancers that it occupies in mESCs and EpiLCs (**Supplementary Figure 6F–H**) near *Lhx5*, *Six3*, and *Sox1* (30, 111). Thus, a decline in TET1 expression upon pluripotency exit is accompanied by a promoter-dominant redistribution of the remaining TET1-bound fraction. Consistent with TET1 peaks concentrating within promoters in early NPCs, TET1 overlap with pH2A.X at active promoters and macroH2A.2 at bivalent promoters also increased (**Figures 3E–F and 4A–B**). Thus, promoter-biased redistribution of TET1 may contribute to the increased recovery of H2A variants in TET1 chromatin complexes from early NPCs. We integrated Day 2 TET1 ChIP-seq and published TET1 KO RNA-seq (31) to determine the regulatory consequences of TET1 loss at bound promoters in early NPCs (**Supplementary Figure 7A–B**). TET1-bound DEGs were upregulated 80% (48/60) of the time after TET1 loss (**Figure 4C**), indicating that TET1 retains its predominantly repressive function after pluripotency exit. These TET1-regulated genes did not include established components of the early NPC transcriptional program, leading us to ask whether TET1 repressed alternative gene programs in early NPCs.

**Figure 4.**
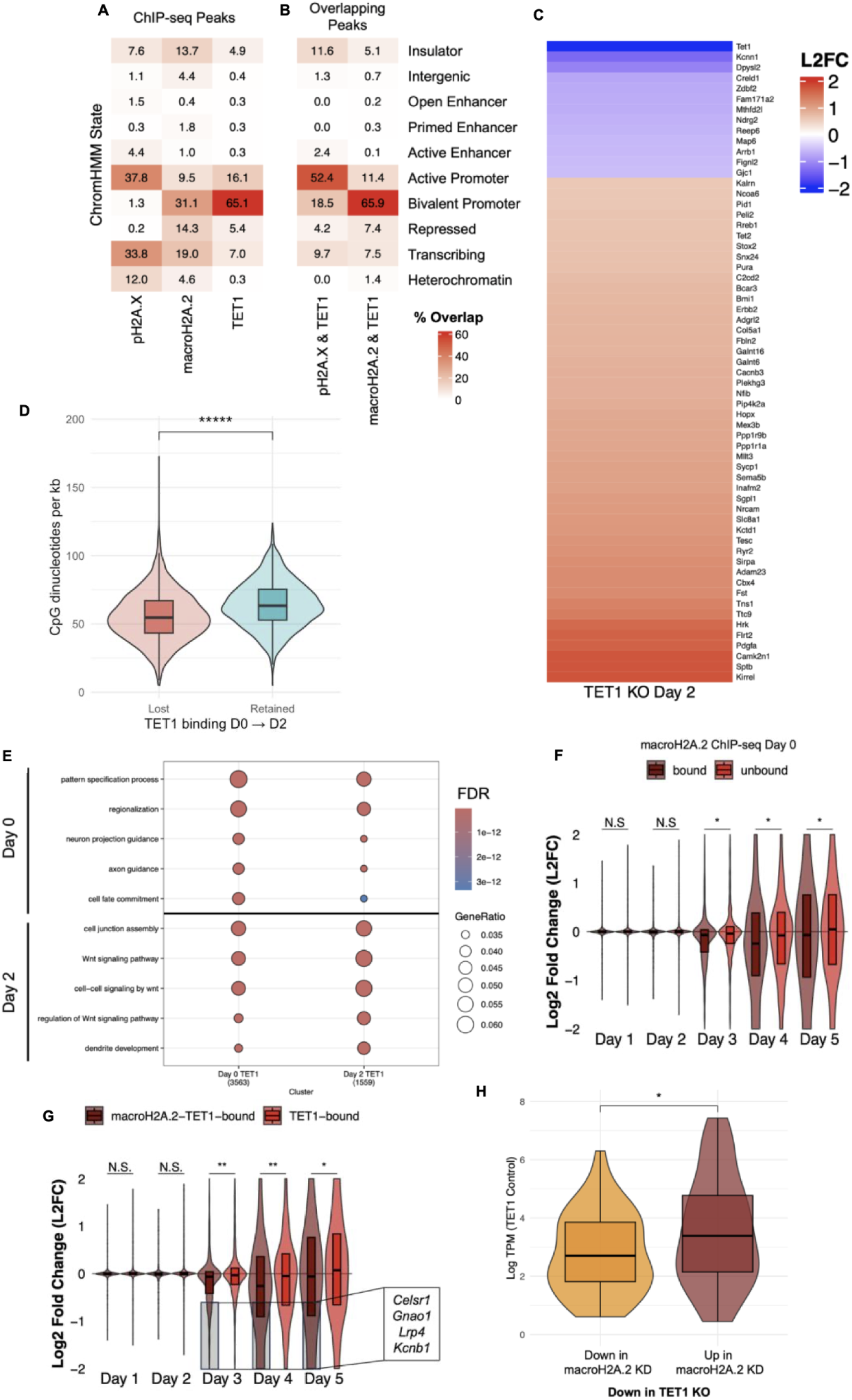
macroH2A.2 mediates TET1-dependent activation of bivalent genes during NPC differentiation. **A.** Log2 FC expression in TET1 KO versus WT at Day 2 of NPC differentiation timecourse (van der Veer et al., 2023) and Day 2 TET1 ChIP-seq targets demonstrates a general pattern of gene activation upon TET1 KO. High-confidence (absolute log fold change > 1.5, FDR < 0.1) differentially expressed genes arranged in ascending order of Log2 FC. **B-C.** ChromHMM Heatmaps of (**B**) ChIP-seq Peaks and (**C**) overlapping ChIP-seq Peaks at genomic regulatory elements in ESCs for pH2A.X, macroH2A.2, and Day 2 of NPC differentiation for TET1 shows Day 2 TET1 enriches in active and bivalent promoters with pH2A.X and macroH2A.2, respectively. Overlap with ChromHMM states is normalised such that each ChIP- seq Peak set sums to 100% across ChromHMM states. **D.** CpG density at bivalent promoters is lower at promoters that lose TET1 binding at Day 2. CpG per kb was quantified from the promoter sequence. A two-sided Wilcoxon rank-sum test was performed with a resulting p-value < 2.2e-16. **E.** Gene ontology analysis of Day 0 TET1-bound bivalent genes versus Day 2 TET1-bound bivalent genes. Day 2 TET1 binding is enriched within bivalent promoters of genes involved in cell-cell contacts, Wnt signaling, and dendrite development. The top 5 terms for each gene list are shown as indicated on the left. An over-representation analysis (one-sided Fisher’s exact test) was performed with a cutoff of FDR < 0.1. GeneRatio is the fraction of queried genes in the gene set. **F.** Log2 fold changes in gene expression (TET1 KO, Day n versus WT, Day n) for DEGs in at least one KO and WT sample during an NPC differentiation RNA-seq timecourse (van der Veer et al., 2023). macroH2A.2-bound at Day 0 (n = 547); macroH2A.2-unbound at Day 2 (n = 1666). A two-sided t-test was performed. * designates an FDR < 0.01. N.S. indicates not statistically significant. **G.** Data represented as in **F**. macroH2A.2-TET1-bound at Day 0 (n = 456); only TET1-bound at Day 0 (n = 1099). A two-sided t-test was performed as in **F**. Each additional asterisk represents an order of magnitude lower FDR. Important genes that are downregulated in the TET1 KO and discussed in the text are indicated in the textbox. **H.** Gene downregulation in NPCs upon TET1 KO predicts lower expression levels when macroH2A.2 KD also leads to downregulation in mESCs. Gene-level log TPM at co-bound genes in WT mESCs (van der Veer et al., 2023). A two-sided t-test was performed. The asterisk indicates a p-value < 0.05.

We first examined loci representing different classes of TET1-mediated dynamic repression between pluripotency and early NPCs. *Mllt3*, a developmental gene with a bivalent promoter in mESCs implicated in retinoic acid receptor A-dependent acquisition of caudal neural fate (112), was bound and repressed by TET1 in pluripotent cells and early NPCs (**Supplementary Figure 7C–D**). Nevertheless, TET1 loss impaired its subsequent activation, suggesting an indirect requirement for TET1 in activating *Mllt3* later in differentiation. *Myo5b*, a developmental gene expressed in neural development with a bivalent promoter in mESCs (113), was bound and repressed by TET1 during pluripotency; however, its subsequent activation during NPC differentiation was unaffected by TET1 loss (**Supplementary Figure 7D**). *Tcea3*, a pluripotency gene that activates the Nodal antagonist *Lefty1* to inhibit mesendoderm development (77), was bound and repressed by TET1 only in early NPCs (**Supplementary Figure 7F**), when it also is downregulated at the protein level (**Figure 1D**). These loci illustrate three modes of TET1 regulation upon transition from pluripotency to early NPCs: an indirect requirement for subsequent activation of the NPC gene *Mllt3*, release from direct repression at the neural gene *Myo5b*, and newly acquired repression of the pluripotency gene *Tcea3*.

We next analysed global TET1 binding dynamics. Across two independent NPC differentiation time-course RNA-seq datasets, loss of TET1 binding was enriched at bivalent genes specifically expressed in early NPCs (two-sided Fisher’s exact tests; this work: OR = 1.46, *P* = 0.0086; van der Veer et al., 2023: OR = 1.38, *P* = 0.021). Strikingly, TET1 retention was strongly associated with higher CpG density within bivalent promoters (**Figure 4D**), consistent with TET1 preferentially binding specifically within CpG dense regions due to affinities independently conferred by its CXXC domain and a downstream region including the CR domain and catalytic core (114). Predictably, bivalent genes were significantly upregulated upon NPC differentiation compared to active pluripotency genes and were most significantly enriched for neurogenesis and pattern-specification Gene Ontology (GO) terms (**Supplementary Figure 8A–B**). TET1-bound bivalent genes in pluripotent cells showed similar functional enrichments to all bivalent genes (**Supplementary Figure 8B; Figure 4E**). In early NPCs, however, TET1-bound bivalent genes were preferentially associated with alternative fates (e.g. Wnt signaling) or later maturation (e.g. dendrite development), whereas terms consistent with NPC differentiation (e.g. axon guidance and cell-fate commitment) were less represented than during pluripotency (**Figure 4E**). For example, *Kalrn* and *Erbb2*, regulators of spine and dendrite morphogenesis and radial glial differentiation, respectively, remained TET1-bound and repressed at both time points and were upregulated only in late NPCs (31, 115, 116). In contrast, *Bmp7* is activated in early NPCs and was repressed by TET1 during pluripotency but not in early NPCs (117). Pluripotency-specific TET1-repressed genes also included Wnt antagonists *Sfrp1* and *Tbx3*; *Sfrp1* represses mesoderm formation and increased during *wt* NPC differentiation, whereas *Tbx3* represses primitive streak formation and declined (118, 119). In early NPCs, retained TET1 targets included *Bcl9l*, a core component of the WNT transcriptional complex (120). Together, these findings support a model in which TET1 broadly represses bivalent developmental genes in pluripotent cells, then selectively disengages from relatively CpG-poor promoters after pluripotency exit to permit early neuroectodermal gene activation while continuing to restrain later neural and alternative-lineage programs.

### MacroH2A.2 modulates bivalent neuroectodermal genes dependent on TET1 for later activation

We next asked what distinguishes bivalent genes in pluripotent cells that require TET1 for activation during NPC differentiation. TET1 occupancy in mESCs or early NPCs was not predictive on its own (**Supplementary Figure 8C–D**). Instead, TET1-dependent activation was associated with macroH2A.2 occupancy (**Figure 4F–G**). Co-bound genes downregulated after TET1 loss in NPCs (Days 3–5) and after macroH2A.2 depletion in mESCs included the neuroectodermal regulators *Celsr1*, *Gnao1*, *Lrp4*, and *Kcnb1* (121–125). Notably, both *Celsr1* and *Tet1* are required for anterior neuropore closure (24, 123). Thus, macroH2A.2 occupancy identifies a subset of TET1-bound bivalent neuroectodermal genes whose subsequent activation depends on TET1.

We then examined how macroH2A.2 regulates this broader class before differentiation. We performed GO analysis of bivalent genes regulated by macroH2A.2 in mESCs that were induced in a TET1-dependent fashion in NPCs (**Supplementary Figure 9**). These genes were enriched for neuroectodermal processes, including synapse assembly and brain development. MacroH2A.2 depletion could either increase or decrease their expression in mESCs, indicating that macroH2A.2 does not uniformly activate neuroectodermal genes but rather modulates them according to their pluripotent-state context. Importantly, many macroH2A.2-repressed genes dependent on TET1 for activation in NPCs were bound but not detectably regulated by TET1 in mESCs. These genes included neural and neuroectodermal regulators such as *Efnb2*, *Kdm2b*, *Rfx3*, *Zic2*, *Six3*, *Sox1*, and *Sox11* (126–132). MacroH2A.2-repressed genes were more highly expressed than macroH2A.2-activated genes within the subset co-bound by TET1 (**Figure 4H**), consistent with the broader regulatory pattern for macroH2A.2 (**Figure 3H**). Yet, there were no significant differences in expression during NPC differentiation between the macroH2A.2-repressed and activated groups. Together, these findings support a temporal model in which macroH2A.2 calibrates expression of poised neuroectodermal bivalent genes that are not necessarily regulated by TET1 during pluripotency but that nevertheless require TET1 for later induction.

## DISCUSSION

Through multiomic analyses, we demonstrate that TET1 regulation during the transition from pluripotency to early NPCs involves dynamic reorganization of TET1 chromatin complexes. Our strategy incorporating biochemical, structural and functional data helps explain the marked variation among reported TET1 KO phenotypes (114, 133–136). Our data show that TET1 binds H2A variants (**Figure 2A–E**), consistent with previous evidence that histones bind the homologous TET2 CR domain and recruit it to chromatin (137). We therefore propose that the histone- and DNA-binding activities of the TET1 CR domain mediate the chromatin-localization function previously mapped to an unspecified region downstream of the CXXC domain (91, 114). This broader region also contains TET1’s nuclear localisation signals and interaction domains with the corepressors SIN3A and PRC2 (43, 138, 139). Its absence is associated with the most severe TET1 KO phenotypes (**Figure 5A–B**), whereas KO strategies that retain it produce relatively mild phenotypes (24, 114, 135, 136). Differences in TET1 KO severity may therefore reflect the extent to which each strategy disrupts nuclear-localization signals, corepressor-interaction sites, and CR-domain-mediated chromatin targeting.

**Figure 5.**
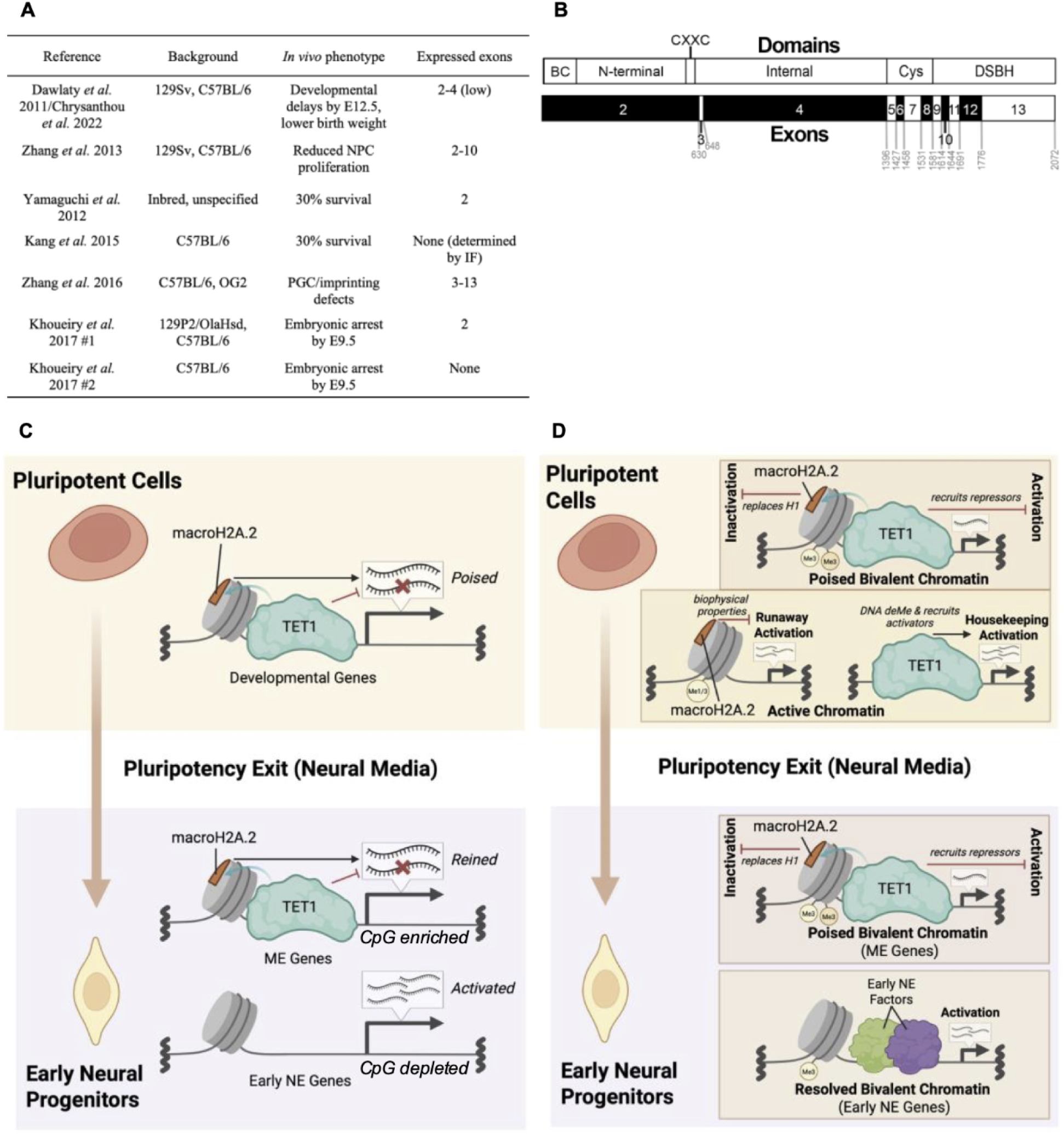
Model of macroH2A.2–TET1 regulation of bivalent genes in pluripotency and NPC differentiation. **A.** Published TET1 knockout strategies and corresponding phenotypes. **B.** Mouse TET1 protein domains aligned to *Tet1* exons. The amino acid corresponding to each exon boundary is indicated in gray. BC = Before CXXC domain; Cys = Cysteine-rich domain; DSBH = double-stranded β helix domain. **C.** Model of macroH2A.2–TET1 functional antagonism at bivalent gene promoters. In mESCs, TET1 represses while macroH2A.2 activates bivalent promoters of lowly expressed, developmental genes. After pluripotency exit at Day 2 of neural progenitor cell differentiation, TET1 is consolidated on macroH2A.2-containing chromatin and represses CpG-enriched promoters of mesoendoderm (ME) but not early NE genes. Created in Biorender.com. **D.** Model as depicted in **C**, with added potential activating and repressive mechanisms (in italics) as described in the main text. Created in Biorender.com.

Our findings support a novel mechanism for selective bivalent gene activation during lineage commitment. We propose a model in which TET1–macroH2A.2 functional antagonism and locus-selective TET1 disengagement partition bivalent genes between activated early-neuroectodermal programs and persistently repressed alternative-lineage or later neural programs. In mESCs, the opposing effects of TET1 and macroH2A.2 support a low-expression yet activation-competent state at bivalent promoters. Upon neuroectodermal commitment, TET1 abundance declines and its binding is broadly lost, particularly from distal regions (**Supplementary Figure 6E**). Nevertheless, early-NPC TET1 binding at a subset of neuroectodermal enhancers primed during pluripotency is associated with enhancer activation (**Supplementary Figure 10**), suggesting that TET1 enhancer regulation is functionally important beyond pluripotency (1, 31). At promoters in early NPCs, TET1 is preferentially retained at CpG-dense loci (**Figure 4D–E**), where it protects the early NPC gene program and further NPC differentiation potential (24, 29, 31, 140, 141). At these loci, TET1 maintains repression of alternative-lineage and late-neural programs (**Figure 5C**), including Wnt-pathway genes such as *Bcl9l*, a component of the WNT transcriptional complex, restricting mesoendodermal differentiation (29, 31, 118, 120).

We propose that this redistribution reflects a locus-selective response to declining TET1 abundance: TET1 occupancy decreases preferentially at lower CpG density, macroH2A.2-bound promoters of genes activated in early NPCs, relieving repression and potentially permitting their activation by early NPC transcription factors. In this model, as TET1 concentration becomes limiting, its binding affinity for CpG sites favors its retention at CpG-rich promoters. This concentration-dependent mechanism provides a biochemical framework for the selective activation or continued repression of genes that are bivalent during pluripotency. Overall, our model explains why TET1 is required after pluripotency exit for bivalency resolution and proper differentiation (6).

We further show that distinct H2A variants mark TET1 activation and repression domains (**Figure 3E–H**). At active promoters, our results may be contextualized by prior findings that TET1 catalytic activity demethylates transcription-factor binding sites, recruits co-activators, and cooperates with pH2A.X to promote transcription and R-loop resolution (1, 29, 96, 97, 142–146). Consistent with this activating function at pluripotency genes, H2A.X phosphorylation supports pluripotent cell self-renewal (86–88). MacroH2A.2 has been proposed to repress highly transcribed genes through linker-mediated inhibition of transcription-factor binding (147, 148). On the other hand, at bivalent promoters, macroH2A.2 competition with the linker histone H1 could oppose the H1-mediated compaction that facilitates PRC1-mediated H2AK119ub deposition and restoration of H3K27me3 after replication, reinforcing developmental-gene repression in pluripotent cells (98, 99, 149–153). By limiting H1-dependent compaction during differentiation, macroH2A.2 may preserve the ability of neuroectodermal bivalent genes to activate as TET1-mediated repression declines (**Figure 4E–G**; **Supplementary Figure 9**). Indeed, the developmental consequences of macroH2A loss include embryonic defects, increased perinatal lethality, and disruption of neuroectodermal gene activation and pluripotency and mesenchymal gene silencing (61, 62, 92, 154–157). Thus, the context-dependent functions of TET1 may operate within a broader epigenetic system governing chromatin plasticity (**Figure 5C–D**). Future studies are needed to further characterize such a system during differentiation and during other cell state transitions where chromatin plasticity is regulated by TET1, H2A.X, and macroH2A, such as somatic reprogramming and cellular rejuvenation (62, 158–165).

The novel histone binding function of TET1 may couple its recruitment with the local chromatin-modification landscape, as demonstrated for the DNMT family (166–169). We show that the TET CR domain is conserved between TET1 and TET2, along with its binding site that recognises H3 tails methylated at K36 (**Figure 2D**), a modification also recognized by PRC2 and DNMT3A (137, 170–172). H3K36me2/3 marks bivalent-domain boundaries, antagonizes H3K27me2/3 deposition, and can impede cellular reprogramming (173–175). During pluripotency, these interactions may form part of a broader regulatory network: TET and DNMT enzymes dynamically oppose one another at accessible, cell-type-specific enhancers, while DNA methylation inhibits PRC2 and bivalent histone marks inhibit DNMT activity (144, 145, 176–179). The differing sensitivities of TET1, PRC2, and DNMTs to these modifications may therefore coordinate their chromatin localization and help create a poised, bistable state at bivalent loci capable of switching between repression and activation in response to developmental cues (144, 172, 177, 180–182). Indeed, recent work highlights that regulation of differentiation by bivalent chromatin depends on not only bivalent marks themselves but also additional factors, such as DPPA2/4 and BEND3, which promote and restrict bivalent-gene activation, respectively (183–186). Differences in signal-responsiveness of regulators may thereby shape the context-specific potential of bivalent chromatin.

## DATA AVAILABILITY

All RNA-seq and ChIP-seq data generated in this study are available at https://www.ncbi.nlm.nih.gov/geo/ with accession number ###.

## SUPPLEMENTARY DATA

Supplementary Data are available at ###.

## ACKNOWLEDGEMENTS

We thank the excellent core facilities at the Cancer Research UK Cambridge Institute and the Babraham Institute for their technical support. RIME and qPLEX-RIME sample processing, mass spectrometry and analysis were performed by Joshua Kent, Kamal Kishore and Clive D’Santos and the Cancer Research UK Cambridge Institute Proteomics Core Facility. RNA sequencing was performed by the Sanger Institute Next-Generation Sequencing Facility. ChIP sequencing was performed by the Cancer Research UK Cambridge Institute Genomics Core. Next-generation Sequencing data processing was performed by the Babraham Institute Bioinformatics Core Facility.

## Author contributions

B.T., A.P., and W.R. designed and conceived the study and interpreted all data. A.P. and W.R. directed and supervised the study. B.T. performed all experiments and bioinformatic analyses. F.S. aided in immunofluorescence imaging and quantification. I.C. provided expertise on TET regulation of pluripotency and the TET1-tagged E14 ESC cell line. B.T., A.P., and W.R. wrote the manuscript. B.T. prepared the figures. All authors read, discussed and approved the manuscript.

## FUNDING

Research in W.R.’s laboratory was supported by the Biotechnology and Biological Sciences Research Council (BB/K010867/1), Wellcome (095645/Z/11/Z) and the European Research Council (ERC) under the European Union’s Horizon 2020 research and innovation programme (EpiCell lineage 882798). B.T.’s doctoral studies at the University of Cambridge were supported by the Jack Kent Cooke Foundation and the Gates Cambridge Trust. A.P. was funded by a Sir Henry Wellcome Fellowship (215912/Z/19/Z).

## Conflict of interest statement

None declared.

## Notes

### Competing Interest Statement

The authors have declared no competing interest.

