## Supplementary for "Dynamic TET1 repression coordinates activation of macroH2A.2-poised bivalent genes during neuroectoderm differentiation"

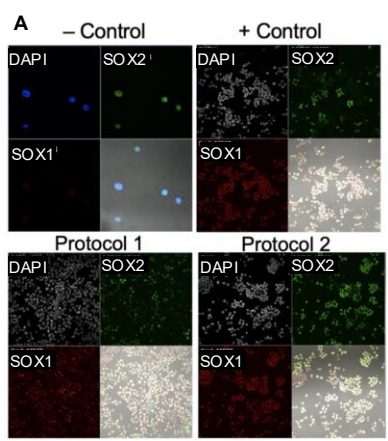

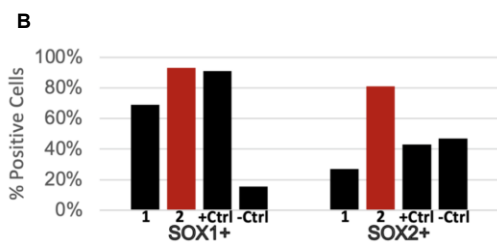

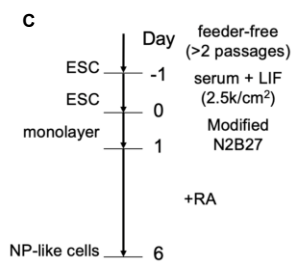

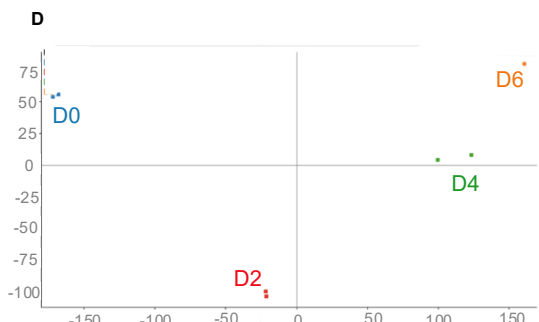

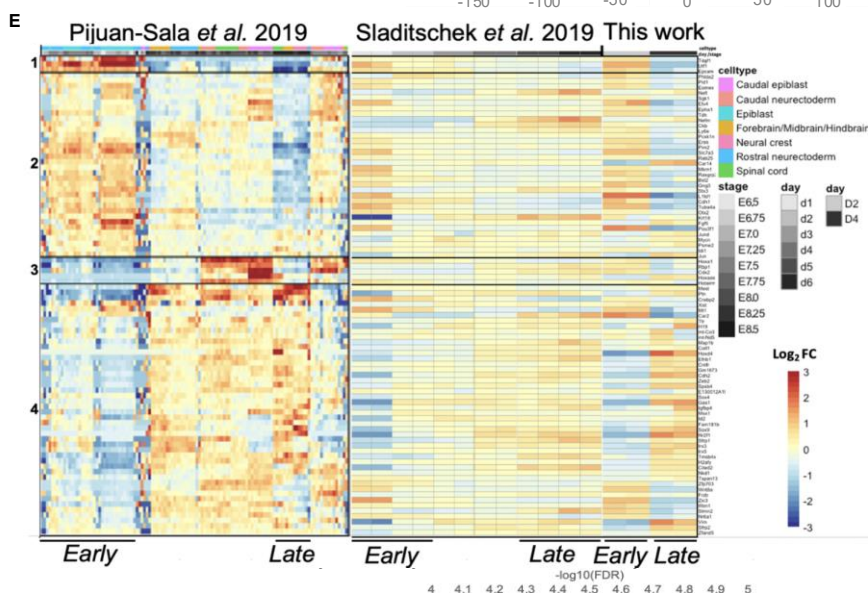

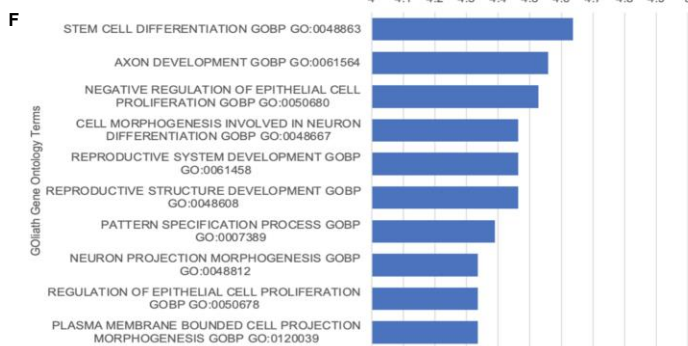

**Supplemental Figure 1.** Quality control and *in vivo* comparison of NPC differentiation.

**A.** Representative IF stainings of NPC differentiation cell cultures from the following conditions: positive control (Day 6, cells plated at 10k/cm<sup>2</sup>, RA-treated), negative control (Day 1, cells plated at 10k/cm<sup>2</sup>, no RA treatment), protocol 1 (Day 6, cells plated at 5k/cm<sup>2</sup>, no RA treatment), protocol 2 (Day 6, cells plated at 2.5k/cm<sup>2</sup>, RA-treated). RA = 1 μM all-*trans*-retinoic acid. **B.** Quantification of IF stainings in **A**. Protocol 2 retains SOX2+ cells more than a previously published modified protocol 1 and control protocol, as measured by immunofluorescence. Percentages calculated from counts of 500 cells per condition. **C.** Schematic of *in vitro* neural progenitor cell differentiation protocol 2, adapted from Ying *et al.* 2003b, used in this study. Cells are replated at Day -1 at 2.5k/cm<sup>2</sup>. NP = neural progenitor; LIF = leukemia inhibitory factor. **D.** PCA of RNA-seq data demonstrates samples separate by timepoint along PC1. **E.** Heatmap of overlapping genes between *in vivo* (Pijuan-Sala *et al.* 2019) and *in vitro* (Sladitschek *et al.* 2019 and this work) neuroectoderm datasets. Early and late groups of samples for each dataset are indicated for comparison. **F.** Top 10 Goliath GO Biological Process Terms for overlapping gene set from **E**. An over-representation analysis (one-sided Fisher's exact test) was performed with a cutoff of FDR < 0.1.

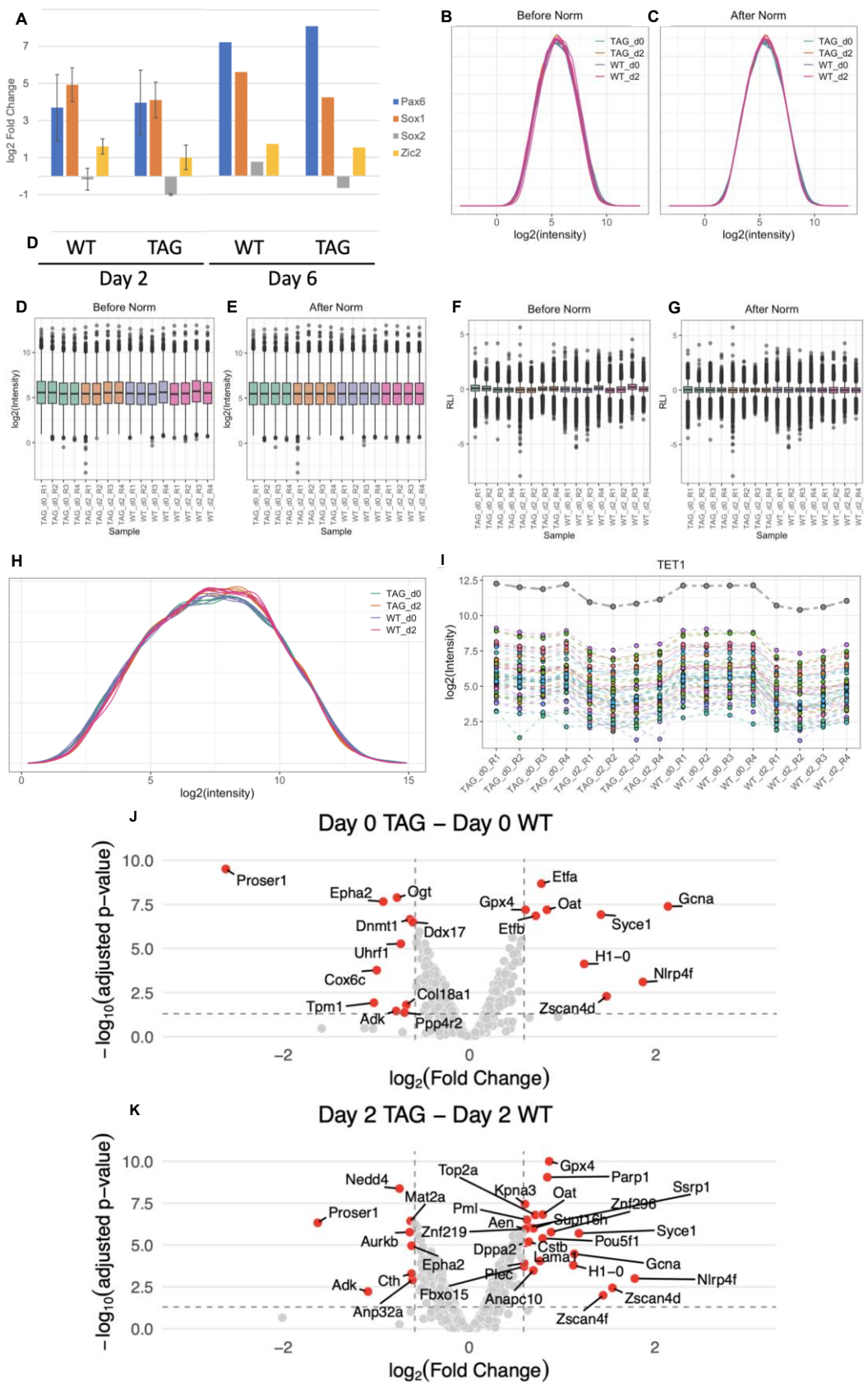

**Supplemental Figure 2.** Quality control of proteomic data from NPC differentiation. **A.** rt-qPCR of cells from Days 0, 2 and 6 of NPC differentiation used in proteome, qPLEX-RIME and ChIP-seq experiments. Days 0 and 2 samples are triplicate while Day 6 samples are duplicate. Days 2 and 6 are compared to Day 0 to calculate $\log_2$ fold change. Error bars represent standard deviation. **B-G.** Full proteome peptide intensities before and after normalization via median scaling. Density plots of **(B)** raw & **(C)** normalized intensities. Boxplots of **(D)** raw & **(E)** normalized intensities. Relative intensity plots of **(F)** raw & **(G)** normalized intensities. **H.** Density plot of log-transformed protein intensities. Protein intensities are summations of peptide intensities. **I.** TET1 protein (gray) is a summation of TET1 peptide (colored) intensities from proteome samples. **J-K.** Volcano plots across replicates of differentially expressed proteins between conditions. **I.** Day 0 TAG versus WT. **J.** Day 2 TAG versus WT. A limma-based statistical analysis was conducted. Differentially expressed proteins (adjusted p-value < 0.05, $|\log \text{ fold-change}| \geq 1.5$) are coloured red and labeled. TAG = TET1-(FLAG)<sub>3</sub> cell line; WT = wildtype cell line.

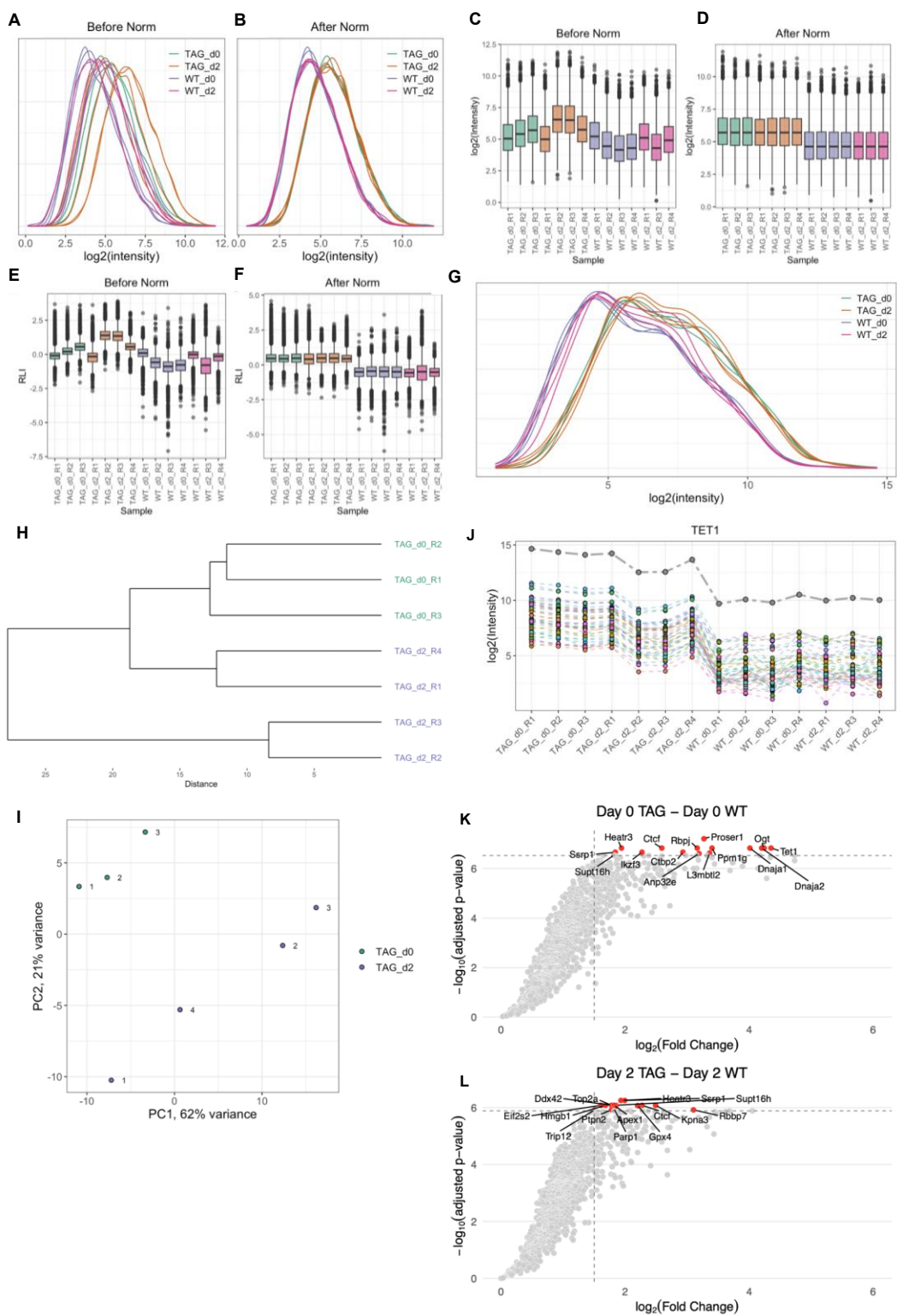

**Supplemental Figure 3.** Quality control of qPLEX-RIME data from NPC differentiation. **A–F.** qPLEX-RIME peptide intensities before and after normalization via median scaling. Density plots of **(A)** raw & **(B)** normalized intensities. Boxplots of **(C)** raw & **(D)** normalized intensities. Relative intensity plots of **(E)** raw & **(F)** normalized intensities. **G.** Density plot of log-transformed qPLEX-RIME protein intensities. Protein intensities are summations of peptide intensities. **H.** Hierarchical clustering of normalized protein intensities using euclidean distance and complete linkage methods. **I.** PCA of normalized protein intensities by condition. **J.** TET1 protein (gray) is a summation of TET1 peptide (colored) intensities from qPLEX-RIME samples. **K–L.** qPLEX-RIME $-\log_{10}$ adjusted p-value (y-axis) versus $\log_2$ fold-change (x-axis) across replicates of differentially enriched proteins between conditions. Day 0 TAG versus WT **(K)**. Day 2 TAG versus WT **(L)**. A limma based statistical analysis was conducted. Top 15 differentially enriched proteins (by adjusted p-value) are coloured red and labeled. TAG = TET1-(FLAG)<sub>3</sub> cell line; WT = wildtype cell line.

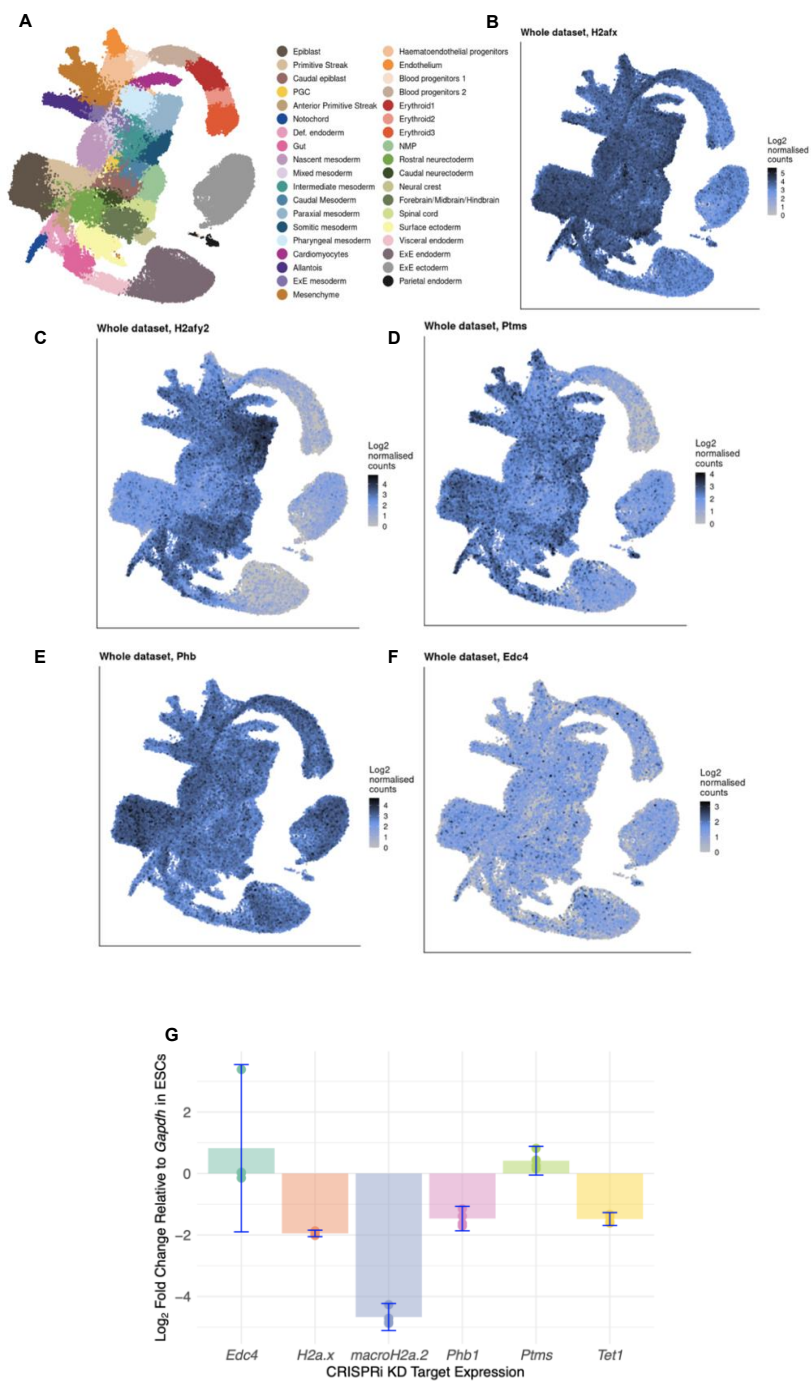

**Supplemental Figure 4.** Testing TET1 qPLEX-RIME hits for physical and functional interactions. **A–F.** 5 qPLEX-RIME-specific hits are expressed in neuroectoderm in the mouse gastrulation atlas. **A.** Gastrulation atlas cell type UMAP. **B–F.** Expression in gastrulation atlas of *H2A.X* (**B**), *macroH2A.2* (**C**), *Ptms* (**D**), *Phb1* (**E**), *Edc4* (**F**). Data from Pijuan-Sala *et al.*, 2019. Plots generated using the atlas Shiny App at marionilab.cruk.cam.ac.uk/MouseGastrulation2018. **G–H.** CRISPRi KD in mESCs (**G**) or NPCs (**H**) of candidate functional interactors with TET1. **G.** rt-qPCR demonstrates efficient CRISPRi KD of *Phb1*, *Tet1*, *macroH2a.2*, and *H2a.x* in mESCs. Experiments performed in quadruplicate with WT mESCs as negative control. Error bars indicate 95% confidence intervals.

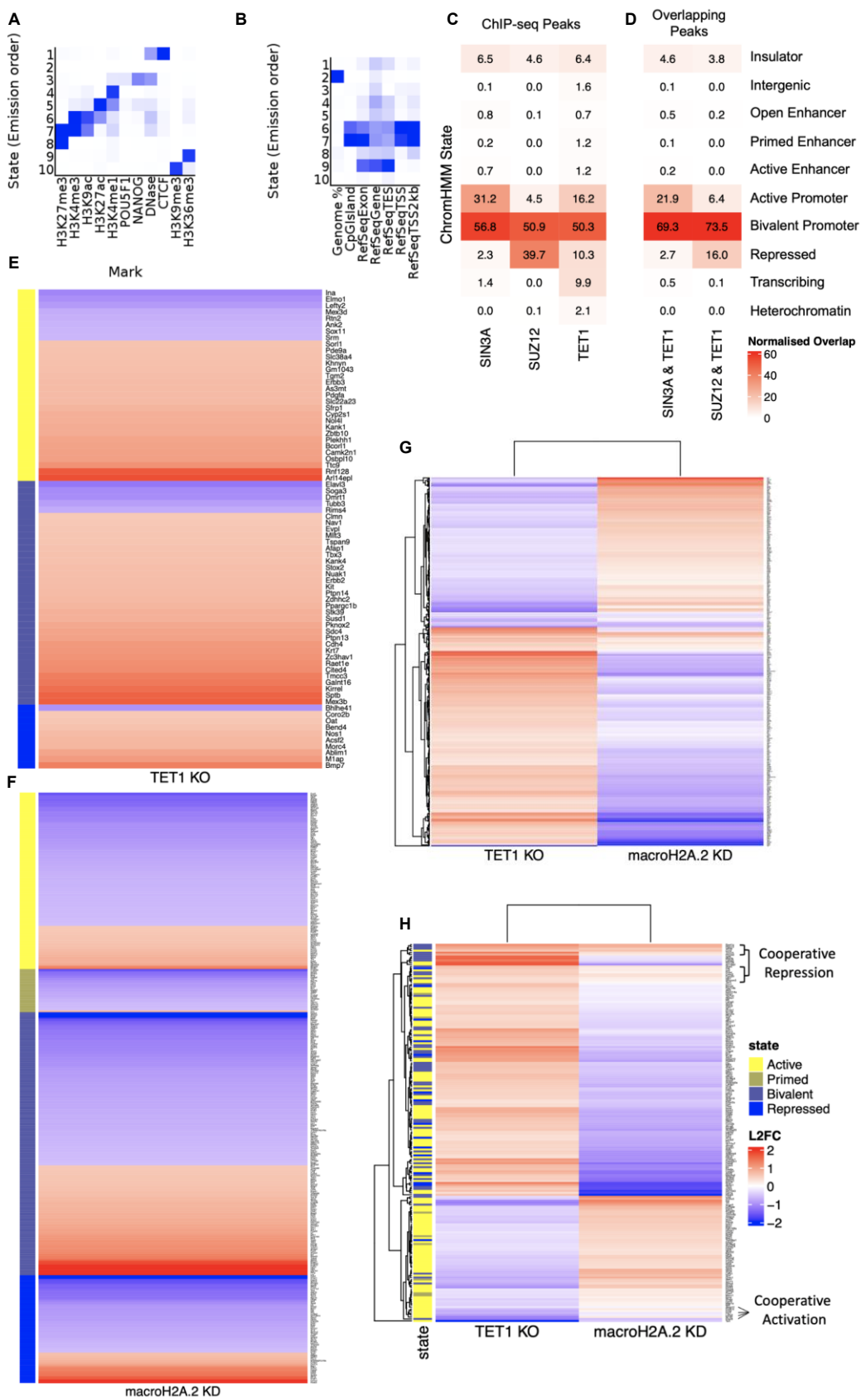

**Supplemental Figure 5.** TET1 regulatory dynamics and partners across ChromHMM states in mESCs. **A.** Heatmap of ENCODE chromatin mark enrichment across ChromHMM states in mESCs. **B.** Heatmap of genome annotation enrichment across ChromHMM states in mESCs. **C–D.** SIN3A, SUZ12 and TET1 ChIP-seq overlap with chromatin states in mESCs. ChromHMM heatmaps of **(C)** ChIP-seq Peaks and **(D)** overlapping ChIP-seq peaks at genomic regulatory elements in ESCs. Overlap with ChromHMM states is normalised such that each ChIP-seq Peak set sums to 100% across ChromHMM states. **E.** TET1 tends to repress genes in mESCs. Log<sub>2</sub> FC expression in TET1 KO versus WT mESCs (van der Veer *et al.*, 2023) at TET1 ChIP-seq targets demonstrates a general pattern of gene activation upon TET1 KO. High-confidence (absolute log<sub>2</sub> fold change > 1.5, FDR < 0.1) differentially expressed genes are grouped by ChromHMM state at their promoters, then arranged in ascending order of log<sub>2</sub> FC. States labelled and expression scaled as in **H.** **F.** macroH2A.2 tends to activate genes in mESCs. Log<sub>2</sub> FC expression in macroH2A.2 KO versus WT mESCs (van der Veer *et al.*, 2023) at macroH2A.2 ChIP-seq targets demonstrates a general tendency toward gene repression upon macroH2A.2 KO. High-confidence differentially expressed genes are grouped by ChromHMM state at their promoters, then arranged in ascending order of log<sub>2</sub> FC. States labelled and expression scaled as in **H.** **G.** TET1 and macroH2A.2 regulate gene expression in opposite directions. Log<sub>2</sub> FC expression of shared DEGs (FDR < 0.1) in TET1 KO and macroH2A.2 KO versus WT mESCs (van der Veer *et al.*, 2023) demonstrates a general pattern of opposite effects of TET1 and macroH2A.2 on gene expression. Genes are clustered by expression pattern. Expression scaled as in **H.** **H.** TET1 and macroH2A.2 regulate gene expression from annotated promoters in opposite directions. Log<sub>2</sub> FC expression of shared DEGs (FDR < 0.1) in TET1 KO and macroH2A.2 KO versus WT mESCs (van der Veer *et al.*, 2023) annotated by promoter ChromHMM state demonstrates a general pattern of opposite effects of TET1 and macroH2A.2 on gene expression. Genes are clustered by expression pattern.

**A** TET1 ChIP-seq Peaks

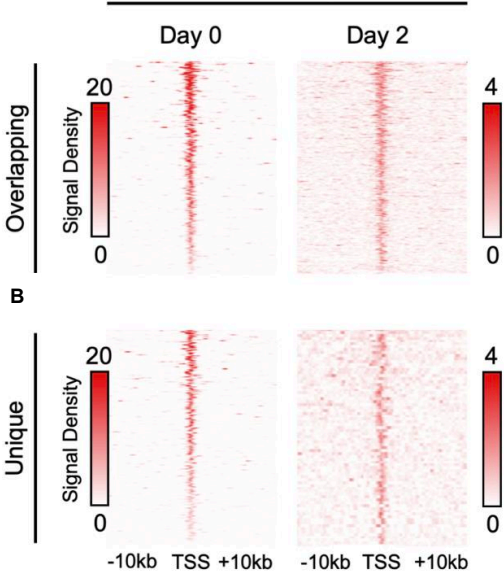

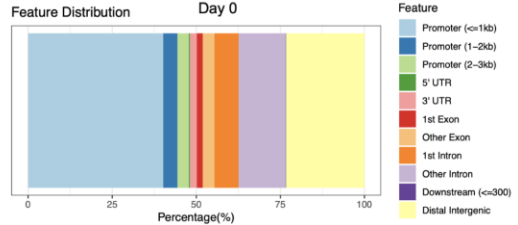

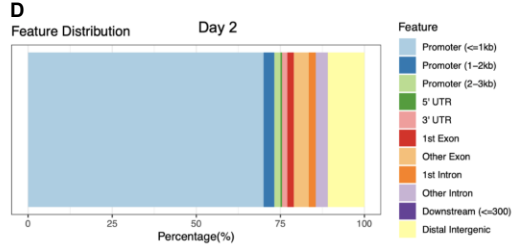

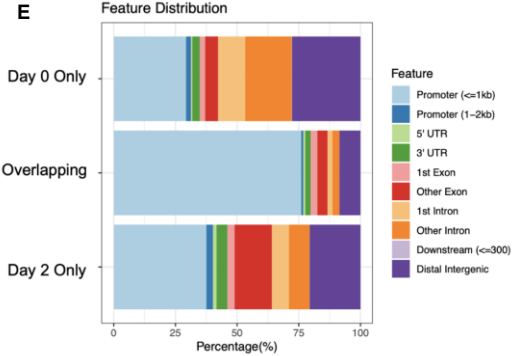

**F**
*Six3* Enhancer
(-133 kb)

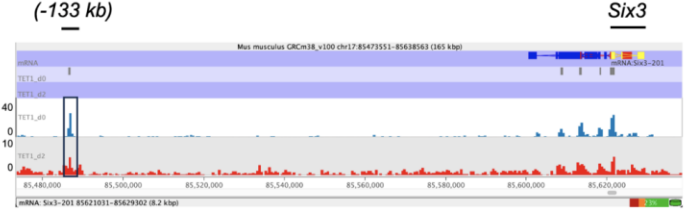

**G** *Sox1* *Sox1 Enhancer*
(+35 kb)

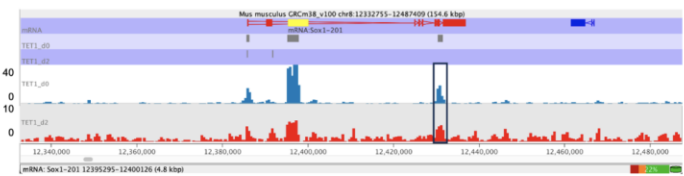

**H** *Lhx5* Enhancer
(-22 kb) *Lhx5*

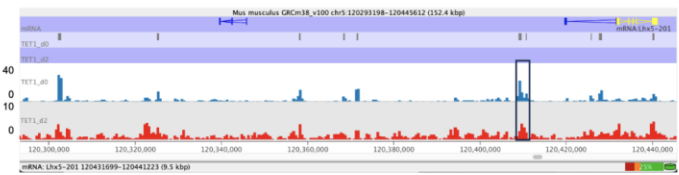

**Supplemental Figure 6.** TET1 binding dynamics in mESCs versus early NPCs (Day 2 of NPC differentiation). **A–B.** Heatmaps of normalised Day 0 (**A**) and Day 2 (**B**) TET1 ChIP-seq TSS-centred read counts. Overlapping peaks between timepoints (upper heatmaps) and unique peaks (lower heatmaps) are shown. **C–D.** Distributions of TET1 ChIP-seq peaks across genomic elements at Day 0 (**C**) and Day 2 (**D**) of NPC differentiation. Promoter distances refer to distances from the transcription start site (TSS). Downstream distance refers to distance from the transcription termination site (TTS). **E.** Distributions of TET1 ChIP-seq peak sets across genomic elements that overlap between timepoints of NPC differentiation. A promoter is defined as the region from 2kb upstream to 0.5 kb downstream of a TSS. **F–H.** Genome browser (Seqmonk) view displaying TET1 ChIP-seq tracks from Day 0 (mESCs) and Day 2 of NPC differentiation show that TET1 binds neural enhancers in serum/LIF mESCs and EpiLCs. **F.** *Six3* enhancer, **G.** *Sox1* enhancer, **H.** *Lhx5* enhancer. Enhancer annotations from Cruz-Molina et al., 2017. TET1 binding data in EpiLCs from Luo et al., 2020.

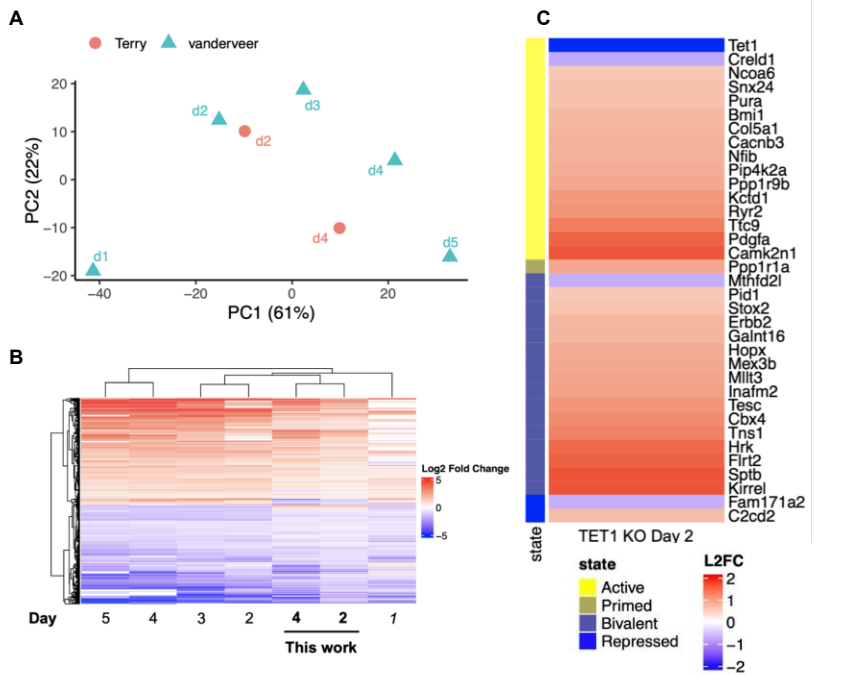

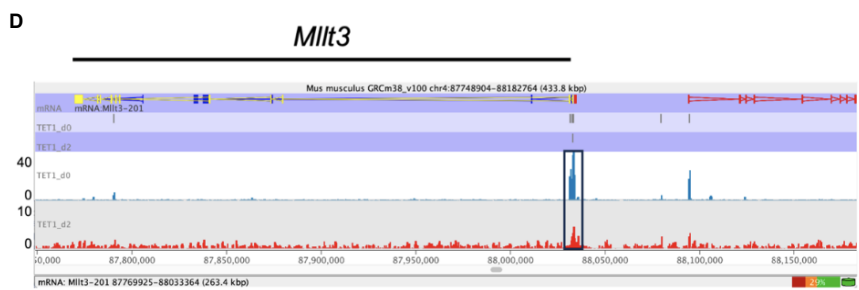

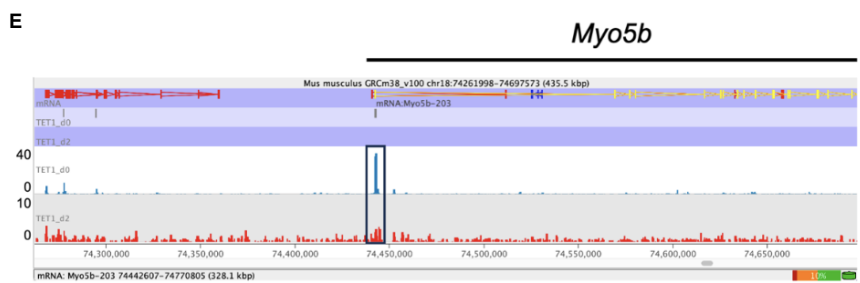

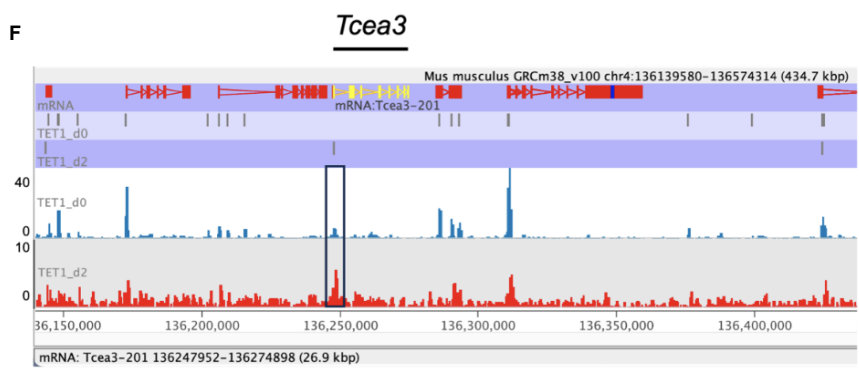

**Supplemental Figure 7. A–B.** Comparison of RNA-seq data from NPC differentiation. **A.** PCA of RNA-seq timepoints from this work (“Terry”) and van der Veer *et al.*, 2023 (“vanderveer”). Top 1000 DEGs were included and limma-based batch effect removal was performed. **B.** Log<sub>2</sub> FC expression of shared DEGs compared to *wt* mESCs (FDR < 0.1) on Day *n* of NPC differentiation timecourses from van der Veer *et al.*, 2023 and this study (in bold). Genes are clustered by expression pattern. **C.** Log<sub>2</sub> FC expression in TET1 KO versus WT at Day 2 of NPC differentiation timecourse (van der Veer *et al.*, 2023) at Day 2 TET1 ChIP-seq targets demonstrates a general pattern of gene activation upon TET1 KO. High-confidence (absolute log fold change > 1.5, FDR < 0.1) differentially expressed genes are grouped by ChromHMM state at their promoters, then arranged in ascending order of Log<sub>2</sub> FC. **D–F.** Genome browser (Seqmonk) view displaying TET1 ChIP-seq tracks from Day 0 and Day 2 of NPC differentiation. **D.** *Myo5b*, **E.** *Mllt3*, **F.** *Tcea3*.

A

Active Bivalent

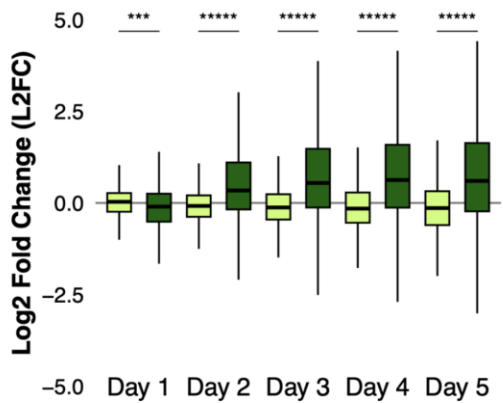

B

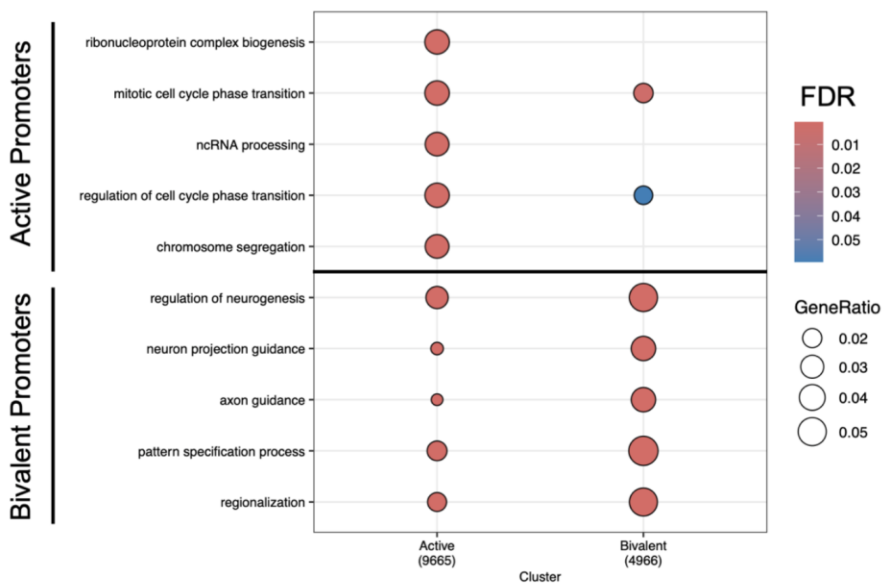

C

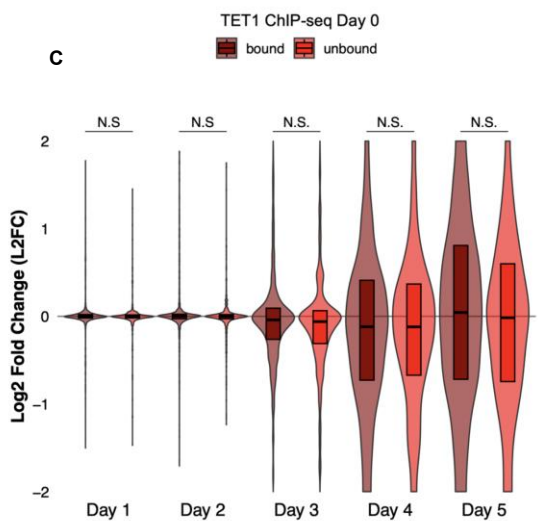

D

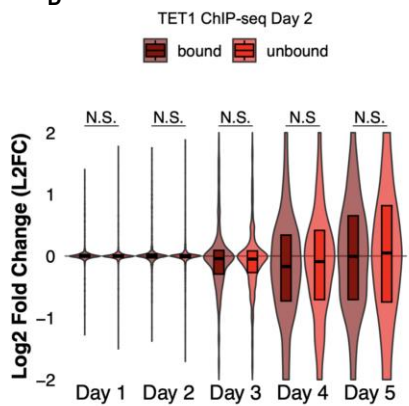

**Supplemental Figure 8.** TET1 binding dynamics alone do not explain bivalent gene regulation during NPC differentiation. **A.** Bivalent genes are activated during NPC differentiation. Boxplots of $\log_2$ fold changes in gene expression at Day $n$ compared to Day 0 of NPC differentiation timecourse (data from van der Veer *et al.*, 2023). A two-sided t-test assuming unequal variance was performed. \*\*\* designates a p-value < 0.0005; \*\*\*\*\* designates a p-value < 1e-100. **B.** Bivalent promoters are enriched in developmental genes. Gene ontology analysis of active versus bivalent genes demonstrate that bivalent genes enrich most significantly for differentiation and neuroectoderm development terms. The top 5 terms for the active and bivalent promoter gene lists are shown as indicated on the left. An over-representation analysis (one-sided Fisher's exact test) was performed with a cutoff of FDR < 0.1. GeneRatio is the fraction of queried genes in the gene set. **C–D.** Bivalent gene expression changes upon TET1 KO do not depend on TET1 binding at Day 0 (**C**) or Day 2 (**D**). $\log_2$ fold changes in gene expression (TET1 KO, Day $n$ versus WT, Day $n$) for DEGs in at least one KO and WT sample during an NPC differentiation RNA-seq timecourse (van der Veer *et al.*, 2023). **C.** TET1-bound at Day 0 (n = 1555); TET1-unbound at Day 0 (n = 658). **D.** TET1-bound at Day 2 (n = 717); TET1-unbound at Day 2 (n = 1496). A two-sided t-test was performed. Significance threshold was set at FDR < 0.01. N.S. indicates not statistically significant.

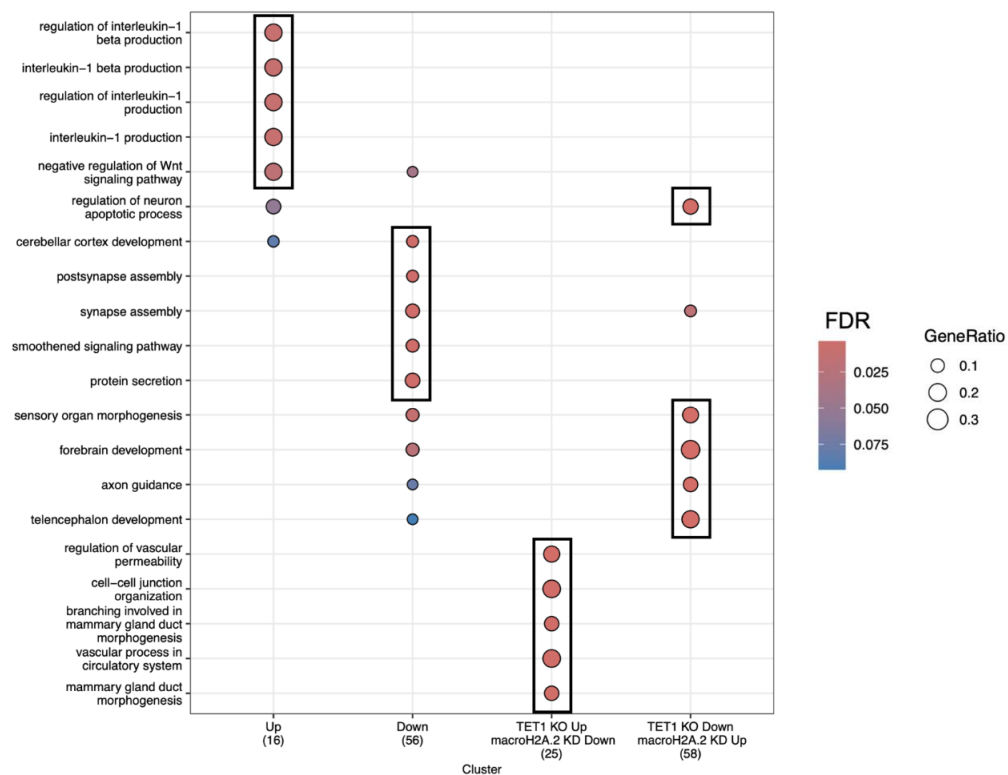

**Supplemental Figure 9.** TET1–macroH2A.2 bivalent gene regulation across NPC differentiation. Gene ontology analysis in NPCs of non-overlapping sets of bivalent DEGs upon TET1 KO bound by TET1 and macroH2A.2 at Day 0. Data from Days 4 and 5 of NPC differentiation from van der Veer *et al.*, 2023). The top 5 enriched terms for each gene list (labeled on the x-axis) are indicated in boxes. An overrepresentation analysis (one-sided Fisher's exact test) was performed with a significance cutoff of $FDR < 0.1$. GeneRatio is the fraction of queried genes in the gene set.

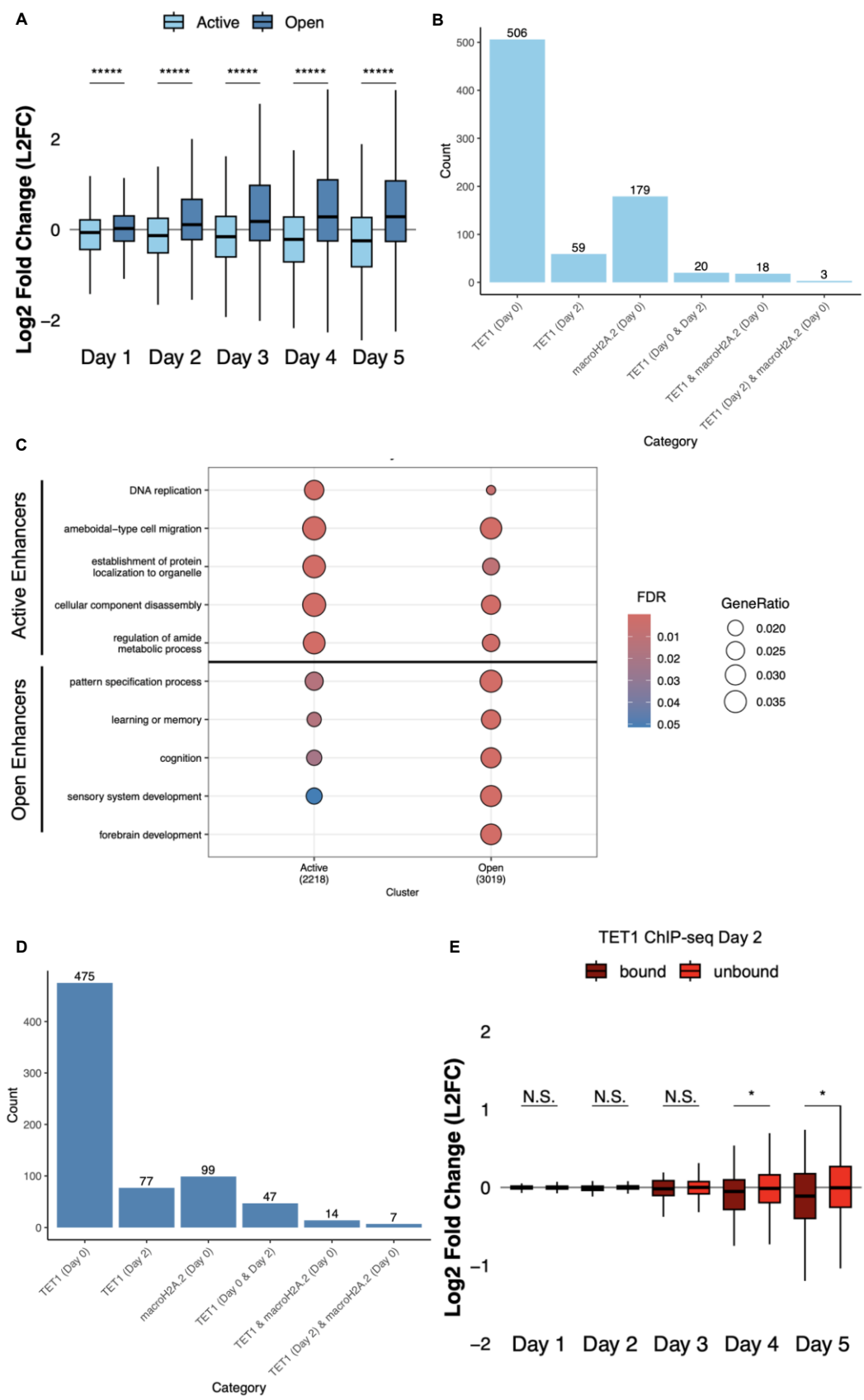

**Supplemental Figure 10.** TET1 activates enhancers open in mESCs during NPC differentiation. **A.** Open enhancers are activated during NPC differentiation. Boxplots of $\log_2$ fold changes in gene expression at Day $n$ compared to Day 0 of NPC differentiation timecourse (data from van der Veer et al., 2023) demonstrate that open enhancer-linked genes ($n = 3118$) are activated during the timecourse while active enhancer-linked genes ($n = 2302$) are inactivated. \*\*\*\*\* designates an $FDR < 2e-16$. **B.** Open enhancers are enriched in genes involved in brain development. Gene ontology analysis of active versus open enhancer-linked gene sets demonstrate that active enhancer-linked genes enriched most significantly for biological process terms related to DNA replication and cell migration, while open enhancer-linked genes enriched most significantly for biological process terms related to forebrain development and pattern specification. The top 5 terms for the active and open enhancer-linked gene lists are shown as indicated. An over-representation analysis (one-sided Fisher's exact test) was performed with a cutoff of $FDR < 0.1$. GeneRatio is the fraction of queried genes in the gene set. **C.** TET1 and macroH2A.2 minimally overlap within active pluripotency enhancers. Upset plot of the number of TET1 Day 0, TET1 Day 2, macroH2A.2 Day 0, and overlapping ChIP-seq peaks between two ChIP-seq datasets within active enhancers. **D.** TET1 and macroH2A.2 minimally overlap within open enhancers. Upset plot of the number of TET1 Day 0, TET1 Day 2, macroH2A.2 Day 0, and overlapping ChIP-seq peaks between two ChIP-seq datasets within open enhancers. **E.** Open enhancer-linked gene expression depends on TET1 binding at Day 2. $\log_2$ fold changes in gene expression (TET1 KO, Day $n$ versus WT, Day $n$) during NPC differentiation RNA-seq timecourse (van der Veer et al., 2023). TET1-bound at Day 2 ($n = 62$); TET1-unbound at Day 2 ($n = 6537$). \* designates a $p$-value $< 0.05$.

Supplementary Table 1

| Gene | Primer sequences 5'→3' (fw/rv) | Source |
| --- | --- | --- |
| Cyclo1B | GACAGACAGCCGGGACAAGC/<br>GGGGATTGACAGGACCCACA | Eckersley-Maslin<br>*et al.*, 2020 |
| Edc4 | AGATGTGCAGATCCGCTTCC/<br>TCGAGACTCTCCAAATGTGAAG | This study |
| H2a.x | TGCTGCCCAAGAAGAGCAG/<br>AAAGGGCCTTTGTGGAGGTG | This study |
| MacroH2a.2 | TTCCCCAGTGGCAGAACTG/<br>ATTCTTCAGCGAGGACGAGC | This study |
| Pax6 | AACAACCTGCCTATGCAACC/<br>ACTTGGACGGGAACTGACAC | Dawlaty *et al.*, 2011 |
| Phb1 | AAGACAGAAAACGGGGCGTC/<br>GGCTGGATTTGTGTGTTCCC | This study |
| Ptms | TTTGTGGTGGAAAAGGCTGAG/<br>GAGAGCTGGTACGCAATGTC | This study |
| Sox1 | GAGTGGAAGGTCATGTCCGAGG/<br>CCTTCTTGAGCAGCGTCTTGGT | Fu *et al.*, 2014<br>(PMID: 24474203) |
| Sox2 | GCGGAGTGGAACTTTTGTCC/<br>GGGAAGCGTGTACTTATCCTTCT | Xiao *et al.*, 2017<br>(PMID: 28737489) |
| Tet1 | TGCACCTACTGCAAGAATCG/<br>AAATTGGCATCACAGCTTCC | This study |
